# *Ex vivo* expansion of skeletal muscle stem cells with a novel small compound inhibitor of eIF2α dephosphorylation

**DOI:** 10.1101/567461

**Authors:** Graham Lean, Matt Halloran, Oceane Mariscal, Solene Jamet, Jean-Phillip Lumb, Colin Crist

## Abstract

**BACKGROUND:** Regeneration of adult tissues requires the activity of rare, mitotically quiescent somatic stem cells. These features are illustrated by the muscle stem cell (MuSC), also known as the satellite cell for its satellite position underneath the basal lamina of the myofiber. Isolation of MuSCs results in their rapid activation of the myogenic program and their subsequent culture ex vivo leads to loss of stem cell regenerative capacity. These shortcomings make MuSCs difficult to study, manipulate and prevent cell based therapies. We have previously shown that muscle stem cells (MuSCs) require tightly regulated protein synthesis through the phosphorylation of eIF2α. Sal003, an analog of salubrinal that blocks eIF2α dephosphorylation, promotes *ex vivo* expansion of MuSCs retaining regenerative capacity after engraftment into the *Dmd*^*mdx*^ mouse model of Duchenne muscular dystrophy.

**METHODS:** Since micromolar concentrations of sal003 (10μM) are required to expand MuSCs *ex vivo*, we undertook a structure relations study to identify novel sal003 analogs with efficacy at lower concentrations. We demonstrate ex vivo expansion of MuSCs isolated from wild-type and mdx mice using new compounds, and use CRISPR/Cas9 genome editing tools to restore dystrophin expression in cultured MuSCs.

**RESULTS:** Here, we have synthesized and screened chemical analogs of sal003 to identify a novel compound promoting the *ex vivo* expansion of MuSCs. The novel compound expands wild-type and *mdx* MuSCs more efficiently than sal003 and also prolongs culture of primary myoblasts from isolated MuSCs.

**CONCLUSIONS:** We identify a novel sal003 analog, C10, with increased potency at lower concentrations. Culture conditions including sal003 or C10 can extend culture of primary myoblasts from isolated MuSCs, which we predict will enable their further study, genetic manipulation and cell based therapies.

## Introduction

Adult tissues with the capacity to regenerate do so by virtue of their somatic stem cells. The regeneration of skeletal muscle is facilitated by normally quiescent muscle stem cells (MuSCs), or “satellite cells”, named for their satellite position beneath the basal lamina of the myofibre[1], which are essential for post-natal repair of skeletal muscle[2–5]. Quiescent MuSCs normally express PAX7 and, in a subset of body muscle, PAX3[6]. In response to injury, MuSCs activate the cell cycle and the expression of members of the myogenic regulatory family of transcription factors (MYF5, MYOD). MuSCs proliferate, differentiate to repair the injured muscle, and self-renew to re-establish the MuSC pool.

The niche plays an important role in maintaining MuSCs and isolation of MuSCs from their *in vivo* niche rapidly influences gene expression [7,8]. Furthermore, *ex vivo* culture of MuSCs results in their rapid entry into the myogenic program and loss of MuSC regenerative capacity[7]. Efforts to understand biophysical and molecular cues that regulate quiescence have been useful to mitigate the loss of MuSC regenerative capacity during *ex vivo* culture[8] [9,10] [11]. We have shown that the phosphorylation of eIF2α is a translational control mechanism regulating MuSC quiescence and self-renewal. Activated MuSCs dephosphorylate eIF2α, increase protein synthesis, and rapidly activate the myogenic program. Pharmacological inhibition of the eIF2α phosphatase Gadd34/PP1 by the small compound sal003[12], a more potent derivative of salubrinal[13], permits the *ex vivo* expansion of MuSCs that retain regenerative capacity, as illustrated by engraftment into the *Dmd*^*mdx*^ mouse model of Duchenne muscular dystrophy[10].

While salubrinal had been identified as a selective inhibitor of Gadd34/PP1[13], the molecular mechanisms underlying its inhibition, and furthermore the mechanisms underlying the increased potency of sal003[12] remain unclear. Previous structure activity relations (SAR) demonstrate that the tricholoromethyl groups of salubrinal are essential for activity, while the quinolone ring terminus is recognized as a key site for modification (R_2_ group) [14,15] Replacement of the R_2_ N-terminal quinolyl group of salubrinal with 4-chlorophenyl led to the more potent sal003 derivative[12].

Here, we ask whether the chemical structure of sal003 can be further modified to improve efficacy, concentrating on the 4-chlorophenyl group that replaced the quinolyl group of the parent salubrinal molecule. We identify a novel analogue of sal003 that expands MuSCs *ex vivo* at lower concentration. We also demonstrate culture conditions containing the novel sal003 analogue permit *ex vivo* expansion, passaging and genome modification of primary adult MuSCs.

## Results

### Identification of novel sal003 analogs maintaining P-eIF2α and promoting MuSC expansion ex vivo

To identify more efficacious analogues of sal003, we synthesized a library of derivatives (Figure 1). Using a modified route previously described[15], a panel of analogues bearing electronically and sterically diverse substituents at R_2_, which corresponds to the region of sal003 that was modified to improve efficacy, were synthesized. Aside from C8, all analogues were comprised of the trichloromethyl (−CCl_3_) core as this functional group had previously been shown to be crucial for activity[14,15]. With this in mind, a variety of thiourea-based aryl groups were incorporated at R_2_ containing both electron-donating and withdrawing groups at differing positions.

**Fig. 1.**
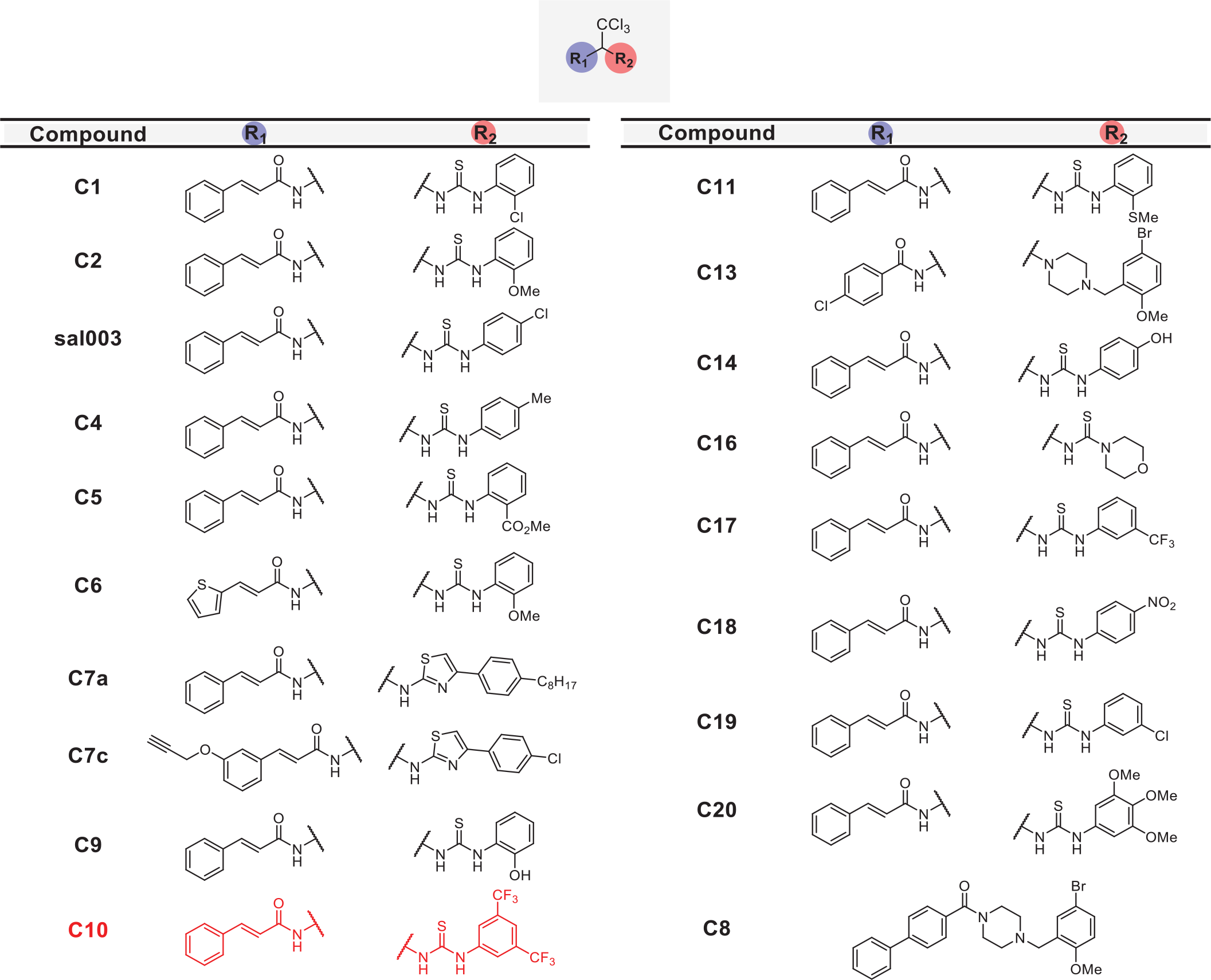
Library of sal003 derivatives.

To screen our panel of novel compounds for activity, we constructed a plasmid vector containing the 5’UTR of *Atf4*, which contain two open reading frames that control *Atf4* mRNA translation in a P-eIF2α dependent manner[16], fused to a firefly luciferase reporter (Fig. 2a). 293 cells were transiently transfected with the *Atf4-fluc* reporter, and treated with sal003 and our panel of novel compounds. We identified four hits that increased P-eIF2α dependent translation of the *Atf4-fluc* reporter above levels achieved by sal003, of which compound 10 (C10) is identified as our lead compound (Fig. 2B). C10 has a 3,5-bis(trifluoromethyl) substituted for the p-chloro on the sal003 N-phenyl group at R_2_ (Fig. 1). Next, we used a quantitative Alpha assay in C2C12 myoblasts to directly determine eIF2α levels after exposure to sal003, C10, a subset of compounds that did not increase translation of *Atf4-luc* reporter (C4, C5, C9 and C11) and 100nM thapsigargin (an ER stress inducer that elevates P-eIF2α). Exposure to C10 resulted in significantly increased levels of P-eIF2α compared to sal003 (Fig. 2c), and these results were confirmed by quantitative western blotting of the P-eIF2α to total eIF2α ratio (Fig. 2d).

**Fig. 2.**
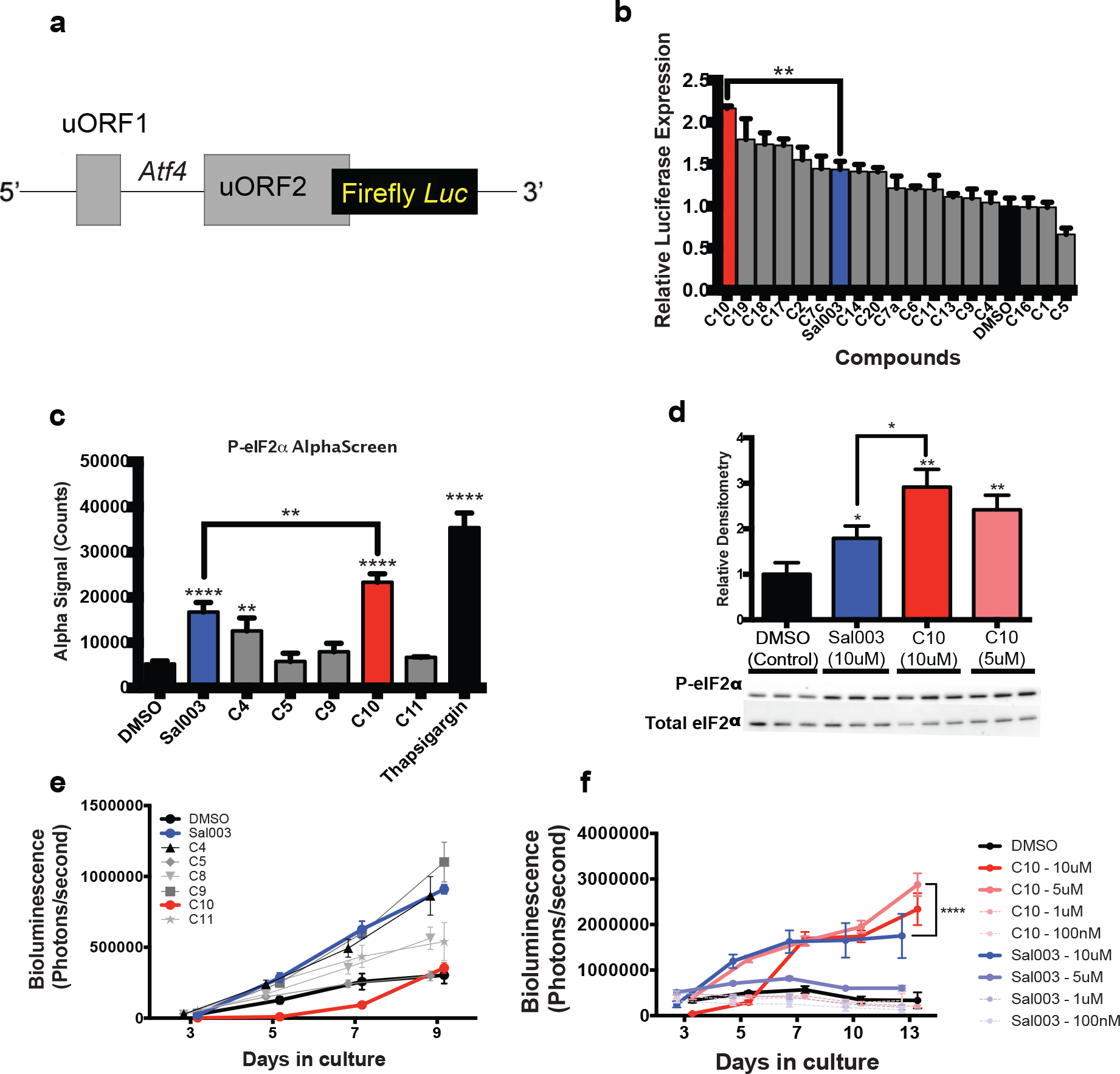
Identification of C10 as a candidate novel small compound inhibitor of eIF2α dephosphorylation. **a** Bioluminescence from MuSCs isolated from *Tg(Cag-luc,-GFP)* adult mice and cultured in the presence of 10μM or DMSO (control). **b** P-eIF2α levels of C2C12 myoblasts after 4 hour culture with 10μM candidate compounds or DMSO (control) using AlphaScreen SureFire eIF2α (Ser-51) assay. **c** Classification of novel compounds based on bioluminescence and P-eIF2a. **d** Bioluminescence from MuSCs isolated from *Tg(Cag-luc,-GFP)* adult mice cultured in the presence of C10, Sal003, at the range of indicated concentration or in DMSO (control). **e** Western blotting against P-eIF2α from cell lysates of C2C12 cells cultured for 4 hours in C10, Sal003 or DMSO (control). All values indicate mean (n≥3) ± SEM.

Next, we monitored the ability of novel compounds to expand *ex vivo* MuSCs isolated from skeletal muscle of adult *Tg(Cag-luc-GFP)* mice, where the expansion of colonies can be observed by bioluminescence (Fig. 2e). Interestingly, one novel compound, C9, which has an o-hydroxyl substituted for the p-chloro on the sal003 N-phenyl group at R_2_ (Fig. 1), caused a rapid increase in bioluminescence (Fig. 2e), without increasing P-eIF2α dependent *Atf4-luc* translation (Fig. 2b) or P-eIF2α levels (Fig. 2c). Meanwhile our lead compound, C10, resulted in poor MuSC expansion at 10μM. We reasoned that excessively high P-eIF2α levels in the presence of 10μM C10 would account for an overall decrease in bioluminescence by promoting lower rates of proliferation, as we have previously shown for sal003[10]. We therefore titrated C10 and sal003 from 10μM to 100nM to show that 5μM C10 potently expands MuSCs (Fig. 2f) at P-eIF2α levels that are more equivalent to 10μM sal003 (Fig. 2d). In contrast, 5μM sal003 does not expand MuSCs *ex vivo* (Fig. 2f).

### Inhibition of eIF2α dephosphorylation by the novel sal003 analog C10 promotes MuSC self-renewal during ex vivo culture

We next examined whether 5μM C10 delays the activation of the myogenic program during *ex vivo* culture. We isolated MuSCs from muscle of adult *Pax3*^*GFP/+*^ mice and cultured them for four days under normal conditions, or in the presence of 5μM C10, and subsequently immunolabelled with antibodies against PAX7, MYOD, and MYOGENIN. Culture in the presence of 5μM C10 over four days resulted in a 2-fold increase in the numbers of PAX7(+); MYOD(−) cells that have not activated the myogenic program and decreased numbers of differentiating PAX7(−), MYOD(+) cells (Fig. 3a, b). Decreased differentiation of MuSCs cultured in the presence of 5μM C10 was also reflected by a decrease in MYOGENIN(+) cells (Fig. 3c, d). Immunoblotting with antibodies against PAX7 and MYOGENIN revealed a 3-fold increase in PAX7 and a 4-fold decrease in MYOGENIN levels after 4 days in culture with 5μM C10 (Fig. 3e, f). In the presence of C10, *MyoD* and *Myogenin* mRNA levels were also reduced. Conversely, *Pax7* mRNA levels did not change (Fig. 3g) to reflect the observed increase in PAX7 protein levels, which is similar to our previous observations with sal003[10].

**Fig. 3.**
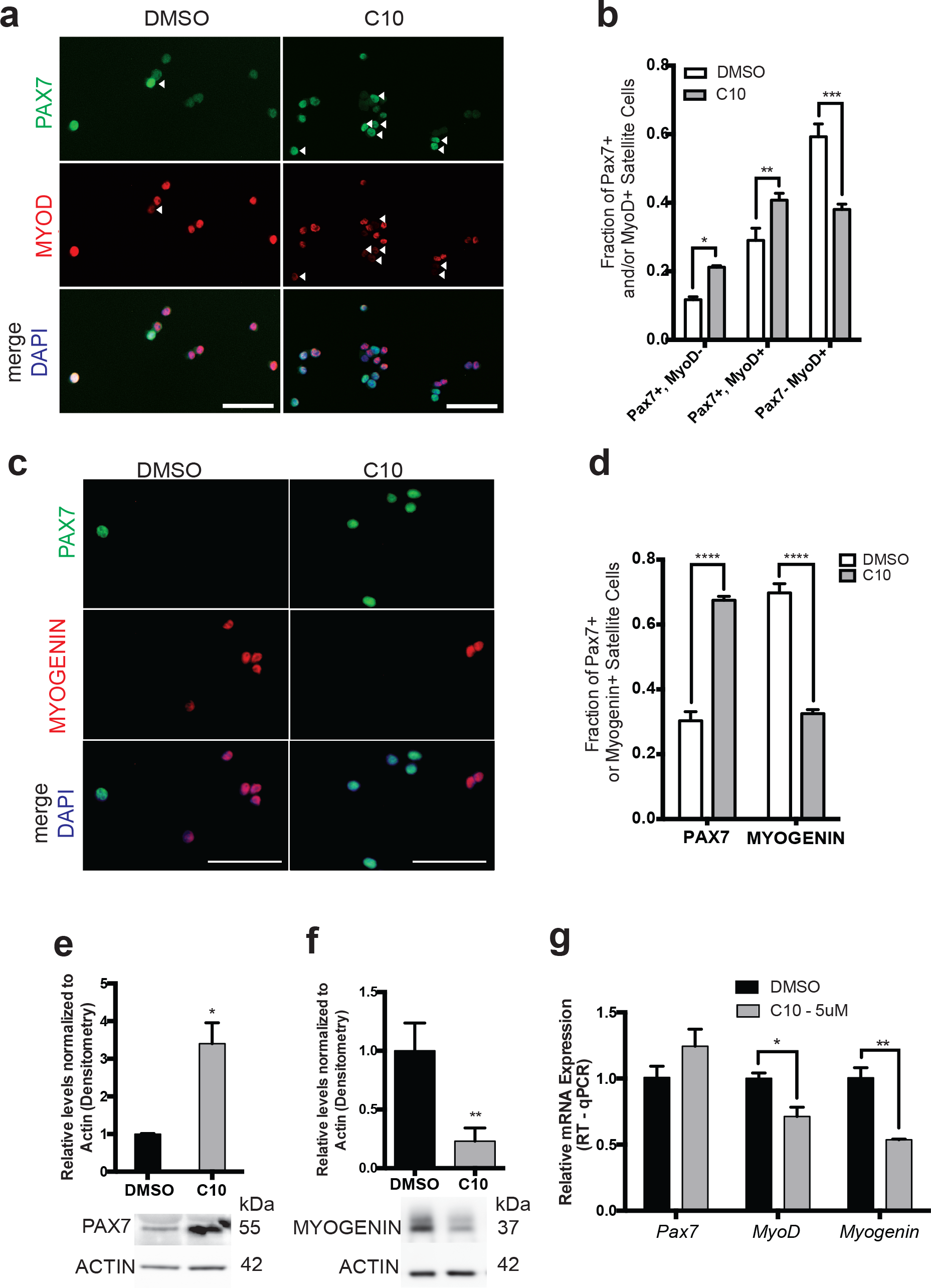
MuSCs cultured in C10 have delayed differentiation and improved self-renewal *ex vivo*. **a** Immunofluorescence with antibodies against Pax7 (Green) and MyoD (Red) on MuSCs isolated from Pax3^GFP/+^ mice after 4 day culture in 5μM C10 or DMSO (control). Quantification of results are to the right of representative images. **b** Immunofluorescence with antibodies against Pax7 (Green) and Myogenin (Red) on MuSCs isolated from Pax3^GFP/+^ mice after 4 day culture in C10 or DMSO (control). Quantification of results are to the right of representative images. **c** Relative mRNA levels of Pax7, MyoD and Myogenin by RT-qPCR after 4 day culture of MuSCs in C10 or DMSO (control). mRNA levels are normalized to *actb* and are relative to DMSO. **d, e** Western blotting with antibodies against Pax7, myogenin and actin of MuSC lysates after 4 day culture in C10 or DMSO (control). Relative densitometry is normalized to actin and presented relative to DMSO (control). Scale bars represent 50μm. All values indicate mean (n≥3) ± SEM.

We next asked whether MuSCs expanded *ex vivo* in the presence of C10 retain their properties to self-renew and regenerate muscle after engraftment into the *Dmd*^*mdx*^ mouse model of Duchenne muscular dystrophy. We isolated MuSCs from muscle of adult *Pax3*^*GFP/+*^ mice and cultured for 4 days in the presence of 5μM C10. After this period of *ex vivo* culture, cells were engrafted into 18Gy irradiated hindlimbs of *Dmd*^*mdx/mdx*^; *Foxn1*^*nu/nu*^ immunodeficient mice. Engraftment of 10 000 C10-treated MuSCs led to higher numbers of DYSTROPHIN(+) myofibres (Fig. 4a, b) and PAX7(+), GFP(+) cells of donor origin (Fig. 4c, d), when compared to 4-day culture of MuSCs in the presence of DMSO.

**Fig. 4.**
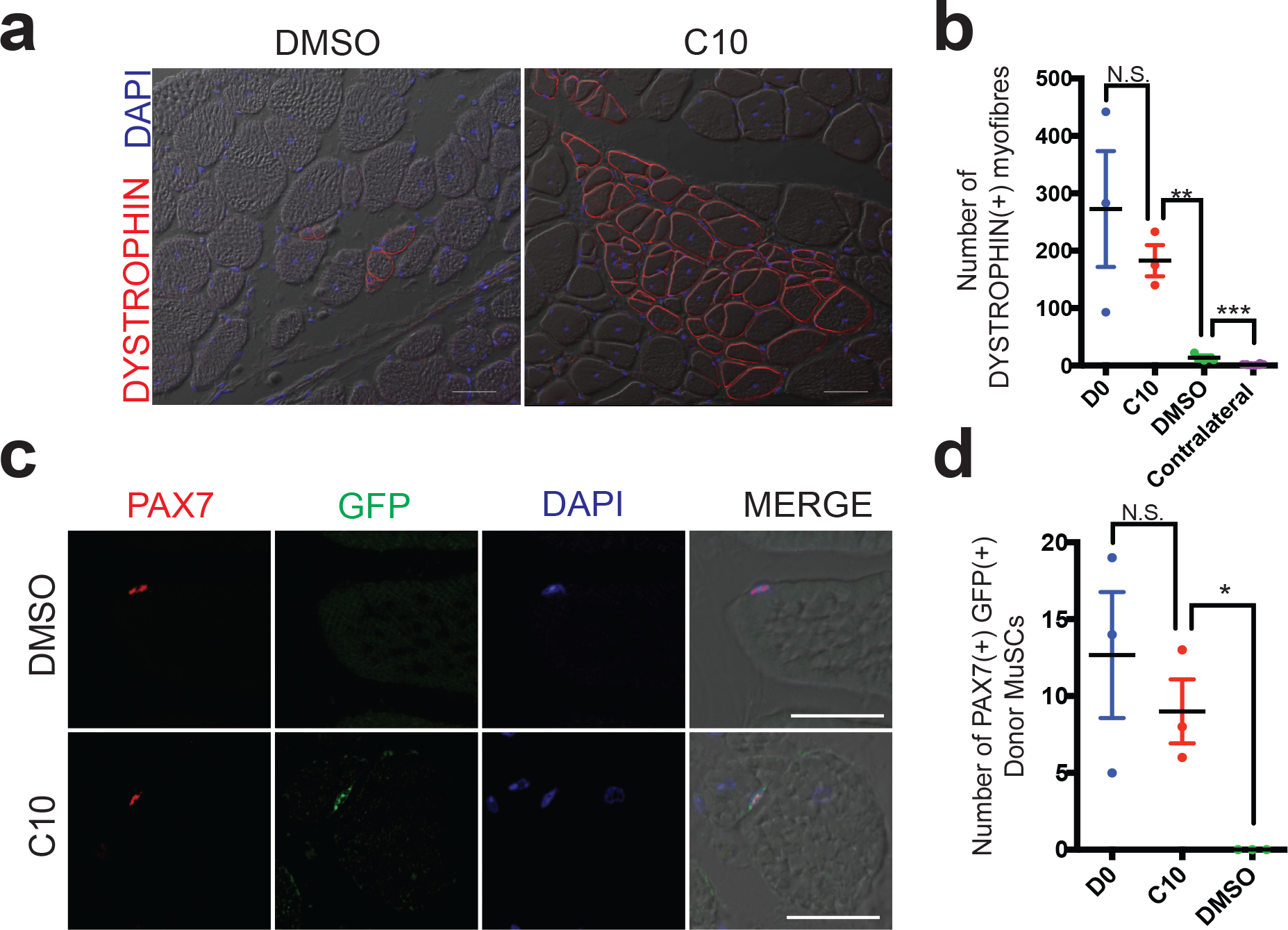
MuSCs maintain their capacity to differentiate and self-renew after 4-day culture in C10. **a** Dystrophin (Red) Immunolabelling of tibialis anterior muscle 21 days after engraftment of 10 000 MuSCs cultured with DMSO (control) or 5μM C10. Scale Bars represents 50μm. **b** Scatter plot of the number of dystrophin(+) muscle fibres per immunolabelled cross-section from (a). **c** Immunolabeling with antibodies against PAX7 (Red) and GFP (Green) on transverse sections of *tibialis anterior* muscle following intramuscular engraftment of MuSCs cultured in DMSO (control) or C10 5μM. Scale bars represent 25μm. **d** Numbers of PAX7(+) GFP(+) donor derived cells per cross-section from (c). All values indicate mean (n≥3) ± SEM.

### Maintaining primary myoblasts from isolated MuSCs in the presence of C10

Primary myoblasts have limited replicative potential, are prone to differentiation, and eventually become senescent in culture. We therefore wanted to determine whether culture conditions including 5μM C10 can extend the utility of primary myoblasts from isolated MuSCs. Isolated MuSCs were seeded on 35mm culture dishes (7500 cells/plate) and cultured in the presence of 5μM C10 or under normal conditions. Every three days, cells were passaged at a density of 1000 cells per 35mm plate. 7500 MuSCs cultured in the presence of 5μM C10 or 10μM sal003 yielded greater than 3 million cells after three passages, compared to an average of 230 000 cells remaining after a maximum of two passages under normal culture conditions (Fig. 5a). After 2 passages, the fraction of PAX7(+) cells is increased by three-fold in the presence of 5μM C10, while the number of MYOGENIN(+) cells is decreased (Fig. 5b, c). After one single passage, extended culture of primary myoblasts in the presence of C10 leads to their eventual differentiation and fusion into myotubes that are larger in diameter (Fig. 5d) and exhibit a higher fusion index (number of DAPI counterstained nuclei per myotube) than normal culture conditions (Fig. 5d, e). Even under these extended culture conditions, the number of PAX7(+) cells that are associated with myotubes are increased 3-fold in the presence of C10 (Fig. 5d, f).

**Fig. 5.**
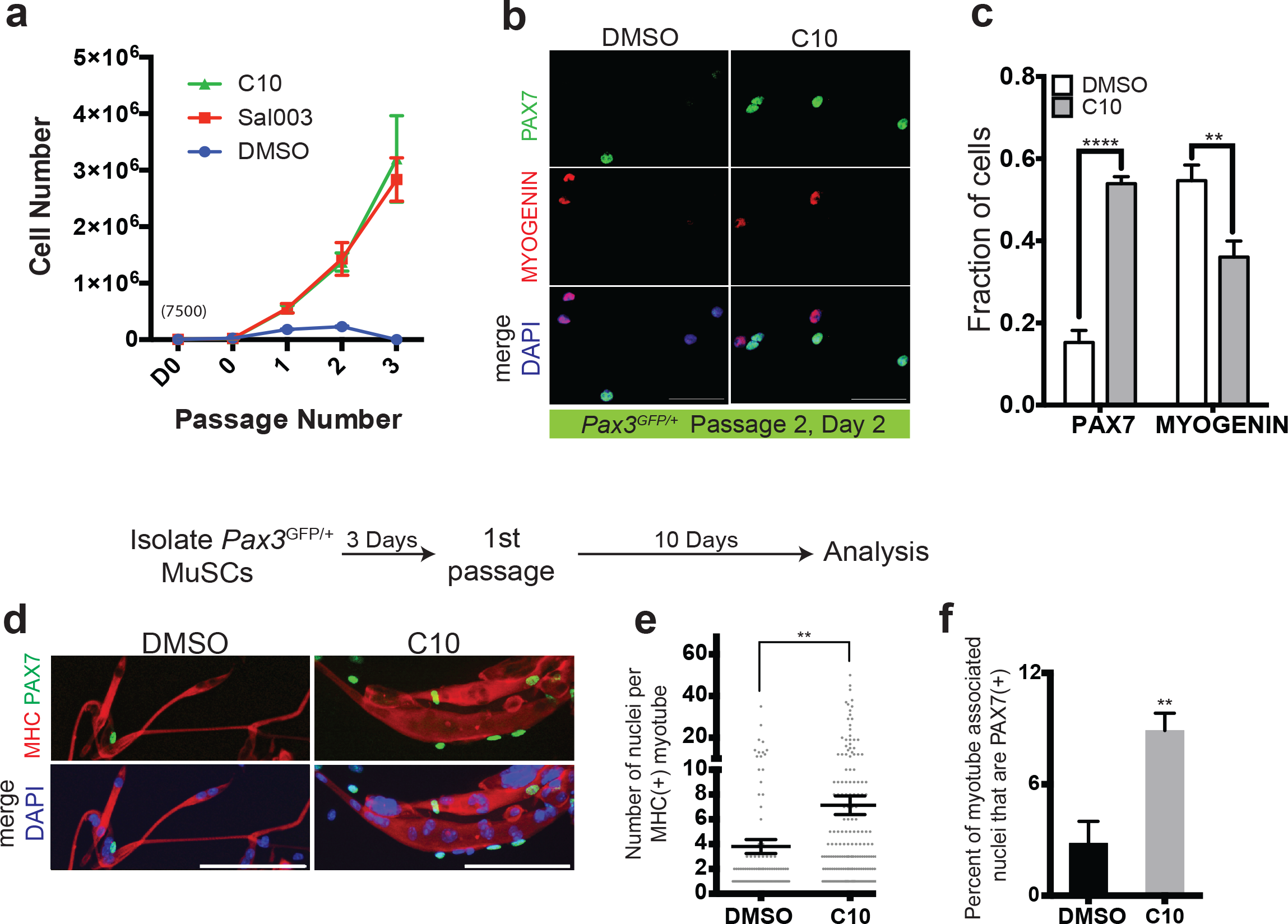
Expansion and passaging of MuSCs in C10. **a** Expansion of MuSCs derived from wild-type mice. MuSCs are plated at an initial density of 7500 cells/plate, passaged at a density of 2000 after 3 days, and then passaged every 3 days at a density of 2000 cells/plate for subsequent passages. **b** Immunolabelling for PAX7 (Green) Myosin Heavy Chain (MHC; Red) of cells cultured for 10 days following passage 1. Scale Bars represent 100μm. **c** Quantification of the number of DAPI+ nuclei per MF20+ myotubes from (b), n≥100 myotubes. **d** Quantification of PAX7(+) nuclei associated with MHC myotubes from (b). **e** Immunolabelling for PAX7 (Green) and MYOGENIN (Red) of wild-type MuSCs after passage 2, cultured with DMSO (control) or 5μM C10. **f** Quantification PAX7(+) or MYOGENIN(+) MuSCs cultured in DMSO (control) or C10 from (e). Scale bars represent 50μm unless indicated otherwise. All values indicate mean (n≥3) ± SEM.

### C10 expands MuSCs isolated from *Dmd*^*mdx*^ mice

We asked whether MuSCs isolated from the *Dmd*^*mdx*^ mouse model of Duchenne muscular dystrophy could be similarly expanded in the presence of C10. We isolated MuSCs from skeletal muscle of *Pax3*^*GFP/+*^; *Dmd*^*mdx*^ (*mdx*) mice and cultured in the presence of 5μM C10 or under normal conditions. Four day culture of *mdx* MuSCs in the presence of C10 yields a two fold increase in numbers of PAX7(+); MYOD(−) cells that have not activated the myogenic program (Fig. 6a, b), a two-fold increase in the total number of PAX7(+) nuclei and a two-fold decrease in MYOGENIN(+) nuclei (Fig. 6c, d).

**Figure 6.**
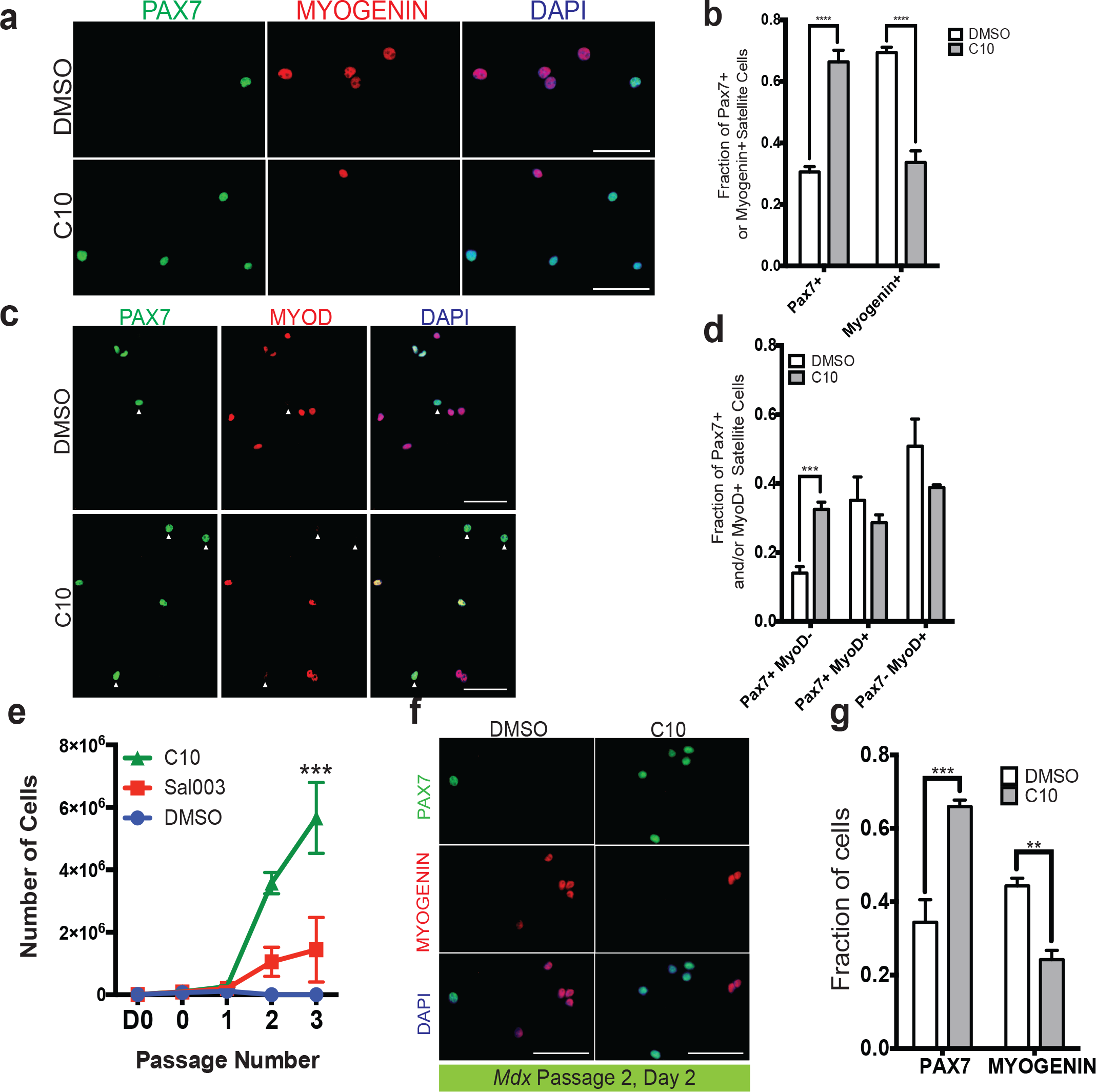
MuSCs isolated from *mdx* mice cultured in C10 have delayed differentiation and improved self-renewal *ex vivo.* **a** Immunofluorescence with antibodies against Pax7 (Green) and MyoD (Red) on MuSCs isolated from *Pax3*^*GFP/+*^; *Dmd*^*mdx*^ mice after 4 day culture with 5μM C10 or DMSO (control). **b** Quantification of results from (a). **c** Immunofluorescence with antibodies against Pax7 (Green) and Myogenin (Red) on MuSCs isolated from *Pax3*^*GFP/+*^; *Dmd*^*mdx*^ mice after 4 day culture with C10 or DMSO (control). **d** Quantification of results from (c). Scale bars represents 50μm. **e** Expansion of MuSCs isolated from muscle of adult *Dmd*^*mdx*^ mice for three passages. MuSCs are plated at an initial density of 7500 cells/35mm plate, passaged at a density of 2000 cells/plate every 2 days. **f** Immunolabelling for PAX7 (Green) and MYOGENIN (Red) of *Pax3*^*GFP/+*^; *Dmd*^*mdx*^ MuSCs after two passages in the presence of DMSO (control) or 5μM C10. **g** Quantification of PAX7(+) or MYOGENIN(+) MuSCs cultured in DMSO (control) or C10 from (f). All values indicate mean (n≥3) ± SEM.

MuSCs were isolated from muscle of adult *Pax3*^*GFP/+*^; *Dmd*^*mdx*^ mice and seeded on 35mm culture dishes (7500 cells/plate) Cells were cultured in the presence of 5μM C10 or under normal conditions and passaged at a density of 2000 cells per 35mm plate every two days. 7500 *mdx* MuSCs maintained in the presence of 5μM C10 yielded on average 5.6 million cells after 3 passages, compared to an average yield of 1.4 million cells in the presence of 10μM sal003 and only 100 000 cells after a maximum of two passages under normal culture conditions (Fig. 6d). After 2 passages, the fraction of PAX7(+) *mdx* cells is increased in the presence of C10, while the number of MYOGENIN(+) *mdx* cells is decreased (Fig. 6e, f).

### Conclusions

Here we have reported a novel compound that expands wild-type and *Dmd*^*mdx*^ MuSCs *ex vivo*, enabling their engraftment *in vivo*, transduction with lentivirus and passaging. Further optimization of sal003/C10 analogs may facilitate the development of cell-based therapies for muscle disease, or potentially overcome defects in endogenous MuSC expansion *in vivo*, which would be expected to delay the progression of muscle disease.

## Materials and Methods

### Compounds

All chemicals and solvents were purchased from Sigma Aldrich, Alfa Aesar, TCI, or Oakwood Chemicals. All solvents were dried and purified using an MBraun MB SPS 800 or Innovative Technology PureSolv MD 7. Unless otherwise stated, reactions were performed in flame-dried glassware under a nitrogen or argon atmosphere. Column chromatography was conducted using 200-400 mesh silica gel from Silicycle. ^1^H-NMR spectra were acquired using Bruker Ascend 500 MHz, Bruker Ascend 400 MHz, and Varian Inova 400 MHz spectrometers. Chemical shifts (δ) are reported in parts per million (ppm) and are calibrated to the residual solvent peak. Coupling constants (*J*) are reported in Hz. Multiplicities are reported using the following abbreviations: s = singlet; d = doublet; t = triplet; q = quartet; m = multiplet (range of multiplet is given). ^13^C-NMR spectra were acquired using Bruker Ascend 125 MHz, Bruker Ascend 100 MHz, and Varian Inova 100 MHz spectrometers. Chemical shifts (δ) are reported in parts per million (ppm) and are calibrated to the residual solvent peak. Analytical thin-layer chromatography was performed on pre-coated 250 mm layer thickness silica gel 60 F254 plates (EMD Chemicals Inc.). Details on the synthesis of sal003 derivatives is available in Supplementary Information.

### Mice

Care and handling of animals were in accordance with the federal Health of Animals Act, as practiced by McGill University and the Lady Davis Institute for Medical Research. All mice are maintained on a C57BL/6 background. For bioluminescence assays, Tg(CAG-Luc-GFP[19] (Jackson Laboratories) were used. For engraftment assays, immunocompromised *Foxn1*^*nu/nu*^; *Dmd*^*mdx4cv/mdx4cv*^ females and *Foxn1*^*nu/nu*^; *Dmd*^*mdx4cv/Y*^ males (Jackson Laboratories) were used. MuSCs were isolated from abdominal and diaphragm muscle of 6-to 8-week old *Pax3*^*GFP/+*^[20], *Pax3*^*GFP/+*^; *Dmd*^*mdx*^ mice using a FACSAriaIII cell sorter as previously described[10]. Alternatively, MuSCs were isolated from the abdominal and diaphragm muscle of 6- to 8-week old *Tg(Cag-Luc,-GFP)* mice via the Miltenyi MACS Satellite Cell Isolation Kit, following by positive selection with Miltenyi Anti-Integrin α7 beads. Isolated MuSCs were cultured in 39% DMEM, 39% F12, 20% FBS, 2% UltroserG, and when indicated, 0.1% DMSO (Control) and 5uM C10, 10uM sal003 or 100nM thapsigargin. MuSC engraftment was performed as previously described [10]. 6- to 8-week-old *Foxn1*^*nu/nu*^; *Dmd*^*mdx*^ mice were used as recipient mice for engraftment assays. Mice were exposed to 18Gy of irradiation to their right hindlimb muscle 24 hours prior to receiving the engraftment. 10 000 cells are suspended in 10μl PBS, loaded into a 10μl Hamilton syringe and introduced in a single injection longitudinally throughout the *tibialis anterior* muscle. Mice were euthanized 21 days after the engraftment and the tibialis anterior muscle was isolated for immunodetection.

### Cell culture assays

C2C12, Hek293 and Hek293T cells were cultured in DMEM with 10% FBS. The *Atf4-luciferase* construct was made by cloning the 5’ UTR of *Atf4* upstream of the firefly luciferase gene in the pGL3 promoter plasmid, as described by Vattem and Wek (2004, PNAS). HEK293 cells were plated in 24-well plates at a density of 25,000 cells/well and incubated overnight. The *Atf4-pGL3* plasmid and pRL-TK renilla luciferase plasmid were co-transfected into these cells using jetPRIME^®^ transfection reagent and incubated overnight. Compounds were reconstituted in DMSO and added to the cells for 16 hours at 10uM final concentration. Cells were then lysed and luciferase expression was measured using the Promega Dual-Luciferase Reporter kit.

P-eIF2α levels were measured using the AlphaScreen^®^ SureFire^®^ eIF2α (P-Ser51) assay kit. C2C12 cells were added to 96-well plates at a density of 5,000 cells/well, and cultured for 24 hours. Compounds were added at a concentration of 10uM for 4 hours and then cells were lysed and lysates were transferred to a 384-well plate. Acceptor beads were incubated with cell lysates at room temperature for 1 hour, and then donor beads were added and incubated overnight at room temperature in darkness. Emission signal was recorded using a PerkinElmer EnVision plate reader.

### Luciferase expansion assay

400 MuSCs from *Tg(Cag-Luc,-GFP)* mice were seeded on a 96-well gelatin-coated plate in MuSC media with 10μM of compounds, or DMSO (control). Bioluminescence (photons/second) was measured at the indicated intervals by replacing 10% of media with MuSC media supplemented with 1.5mg/ml D-Luciferin (Gold Biotechnology) and recorded using an IVIS Spectrum (Perkin Elmer).

### Immunodetection

Immunofluorescence labelling of cultured MuSCs and transverse sections of TA muscle was performed as previously described (Crist et al., 2009, Zismanov et al., 2016). For immunoblotting, cell lysates were prepared as previously described (Crist et al., 2009). ImageJ was used to determine the densitometry from immunoblots. Primary mouse antibodies against PAX7 (DHSB), MYH1E (DHSB, MF20), MYOGENIN (Abcam, EPR4789), b-actin (Sigma, AC-15), P-eIF2a (Abcam, E90), total eIF2a (Cell Signaling, L57A5) and rabbit antibodies MYOD (Abcam), DYSTROPHIN (Pierce), GFP (Life Technologies). Secondary antibodies are Alexa Fluor 488 and 594 anti mouse antibodies, Alexa Fluor 488 and 594 anti rabbit antibodies, and Horseradish peroxidase (HRP) conjugated goat anti-mouse or anti-rabbit antibodies (Jackson Immunoresearch). For immunoblotting, ECL Prime Western Blotting Detection reagents (GE Healthcare) and detected with an ImageQuant LAS 4000 (GE Healthcare).

### RNA analysis

RNA was isolated from cells in culture using TRIzol reagent (Life Technologies) and treated with DNase (Roche). RNA was reverse transcribed using the Superscript III reverse transcriptase (Life Technologies) using oligoDT primers. Primers are *Pax7* Forward (FWD) 5’-CTCAGTGAGTTCGATTAGCCG-3’, Reverse (REV) 5’-AGACGGTTCCCTTTGTCGC-3’; *MyoD* FWD 5’-CCCCGGCGGCAGAATGGCTACG-3’, REV 5’-GGTCTGGGTTCCCTGTTCTGTGT-3’; *MyoG* FWD 5’-CAACCAGGAGGAGCGCGATCTCCG-3’, REV 5’-AGGCGCTGTGGGAGTTGCATTCACT-3’; and *Actb* FWD 5’-AAACATCCCCCAAAGTTCTAC-3’, REV 5’-AAACATCCCCCAAAGTTCTAC-3’.

### Passaging MuSCs

MuSCs were isolated by FACS as previously described and plated on 35mm gelatin-coated dishes at a density of 7500 cells/plate. After 3 days in culture, cells were washed with PBS once and trypsinized with 0.05% trypsin at 37°C for 1 minute. Cells were resuspended in 1ml 10% FBS, F12 media and centrifuged at 600g for 10 minutes at 4°C. Pellets were resuspended in 100μl MuSC media, counted with a haemocytometer and plated at 2000 cells/plate for wild-type and 1000 cells/plate for *Dmd*^*mdx*^. Cells were subsequently passaged every 2 days, following the same protocol.

### Statistical analysis

Graphical analysis is presented as mean ± SEM. At least three independent replicates of each experiment were performed. Unless otherwise indicated, significance was calculated using unpaired Student’s t tests with two-tailed p values: *p < 0.05, **p < 0.01, ***p < 0.001.

