## Supplementary Information for "*Ex vivo* expansion of skeletal muscle stem cells with a novel small compound inhibitor of eIF2α dephosphorylation"

**Experimental Procedures and Data**

### 1. General Experimental

All chemicals and solvents were purchased from Sigma Aldrich, Alfa Aesar, TCI, or Oakwood Chemicals. All solvents were dried and purified using an MBraun MB SPS 800 or Innovative Technology PureSolv MD 7. Unless otherwise stated, reactions were performed in flame-dried glassware under a nitrogen or argon atmosphere. Column chromatography was conducted using 200-400 mesh silica gel from Silicycle. ^1^H-NMR spectra were acquired using Bruker Ascend 500 MHz, Bruker Ascend 400 MHz, and Varian Inova 400 MHz spectrometers. Chemical shifts (δ) are reported in parts per million (ppm) and are calibrated to the residual solvent peak. Coupling constants (*J*) are reported in Hz. Multiplicities are reported using the following abbreviations: s = singlet; d = doublet; t = triplet; q = quartet; m = multiplet (range of multiplet is given). ^13^C-NMR spectra were acquired using Bruker Ascend 125 MHz, Bruker Ascend 100 MHz, and Varian Inova 100 MHz spectrometers. Chemical shifts (δ) are reported in parts per million (ppm) and are calibrated to the residual solvent peak. Analytical thin-layer chromatography was performed on pre-coated 250 mm layer thickness silica gel 60 F254 plates (EMD Chemicals Inc.).

### 2. General Procedures for sal003 Derivatives:

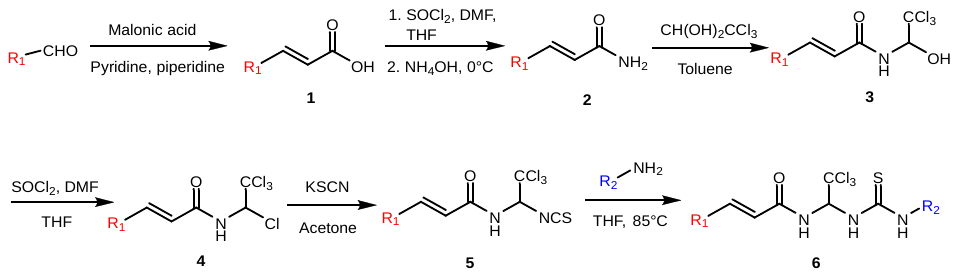

#### a) General Procedure A

A flame-dried round bottom flask the appropriate aromatic aldehyde (1 equiv.) was combined with malonic acid (2 equiv.) and piperidine (0.1 equiv.) in anhydrous pyridine (0.5 M) and heated at 100˚C for 8h. The reaction mixture was cooled to room temperature, quenched with 2M HCl, extracted with DCM, dried with MgSO_4_ and concentrated *in vacuo*. The obtained solid was then washed with water and collected via vacuum filtration to afford **1**.

#### b) General Procedure B

In a flame-dried round bottom flask equipped with a Teflon-coated stir bar, **1** (1 equiv.) was added to a mixture of SOCl_2_ (3 equiv.) and DMF (0.05 equiv.) in anhydrous THF (0.3 M) and heated at reflux for 2h. The reaction mixture was cooled to room temperature, the solvent removed *in vacuo*, and the residue was *carefully* added dropwise to a cooled solution of NH_4_OH (5 equiv.). The obtained solid was then vacuum filtered and washed with water to yield the amide **2**.

#### c) General Procedure C

In a round bottom flask equipped with a Teflon-coated stir bar, **2** (1 equiv.) was combined with chloral hydrate (2 equiv.) in Toluene (0.5 M) and heated at reflux for 12h. The reaction mixture was allowed to cool to room temperature, placed into an ice bath, and the obtained solid was collected via vacuum filtration and washed with cold toluene to yield the chloral derivatives **3**.

#### d) General Procedure D

In a flame-dried round bottom flask equipped with a Teflon-coated stir bar, **3** (1 equiv.) was added to a mixture of SOCl_2_ (3 equiv.) and DMF (0.05 equiv.) in anhydrous THF (0.3 M) and heated at reflux for 2h. The reaction mixture was cooled to room temperature, the solvent removed *in vacuo*, and the obtained solid was washed with cold hexanes and dried under vacuum to yield the chlorinated compounds **4** as solids.

#### e) General Procedure E

In a round bottom flask equipped with a Teflon-coated stir bar, potassium thiocyanate (1 equiv.) was combined with **4** (1 equiv.) and refluxed in acetone (0.5 M) for 1h. The reaction was cooled to room temperature, the white solid was filtered off and the filtrate was concentrated under reduced pressure to afford the isothiocyanates **5**.

#### f) General Procedure F

In a pressure vial equipped with a Teflon-coated stir bar, **5** (1 equiv.) was combined with the appropriate aniline and dissolved in THF (0.2 M). The pressure vial was sealed with the screwcap and heated at 85ºC for 3h. The mixture was cooled to room temperature, the vial was opened, the white solid was vacuum filtered and washed with cold EtOAc to afford the analogs **6*.*** In the case where there was no precipitate formed, the solvent was removed *in vacuo* and the solid was suspended in cold EtOAc and vacuum filtered to obtain **6**.

### 3. Synthesis and Characterization of Compounds in Figure 1

#### a) Synthesis of Starting Materials

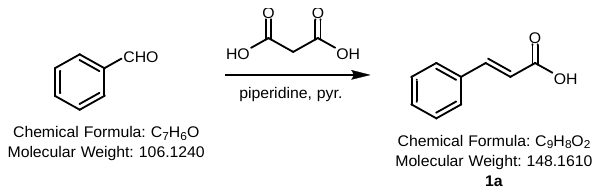

The reaction was carried out according to general procedure A using benzaldehyde (5.31 g, 50.0 mmol, 1 equiv.), malonic acid (10.41 g, 100.0 mmol, 2 equiv.), piperidine (0.49 mL, 5 mmol, 0.1 equiv.), and pyridine (100 mL, 0.5 M). Compound **1a** (6.96 g, 47.0 mmol) was obtained as a white solid in 94% isolated yield.

**^1^H NMR** (300 MHz, Acetone-*d*_6_) δ 10.78 (bs, 1H), 7.75 – 7.65 (m, 3H), 7.48 – 7.40 (m, 3H), 6.55 (d, *J* = 16.1 Hz, 1H) ppm; **^13^C NMR** (75 MHz, Acetone-*d*_6_) δ 167.0, 144.6, 134.6, 130.2, 128.9, 128.1, 118.3 ppm. *Analytical data matches that reported in the literature.^1^*

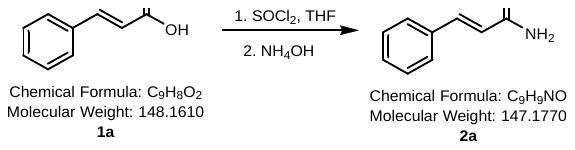

The reaction was carried out according to general procedure B using **1a** (100.0 g. 0.675 mol, 1 equiv.), SOCl_2_ (146.8 mL, 2.025 mol, 3 equiv.), DMF (2.61 mL, 33.75 mmol, 0.05 equiv), THF (675 mL, 1 M), and then NH_4_OH (120 mL, 5 equiv.) to afford **2a** (87.42 g, 0.594 mol) as a white powder in 88% isolated yield.

**^1^H NMR** (300 MHz, Acetone-*d*_6_) δ 7.62 – 7.54 (m, 3H), 7.43 – 7.31 (m, 3H), 7.12 (bs, 1H), 6.75 (d, *J* = 15.8 Hz, 1H), 6.74 (bs, 1H) ppm; **^13^C NMR** (75 MHz, Acetone-*d*_6_) δ 167.0, 140.1, 135.3, 129.4, 128.8, 127.6, 121.7 ppm.

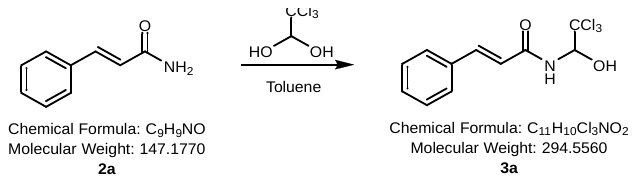

The reaction was carried out according to general procedure C using **2a** (87.42 g, 0.594 mol, 1 equiv.), chloral hydrate (196.49 g, 1.19 mol, 2 equiv.) and toluene (600 mL, 1 M) to obtain **3a** (148.72 g, 0.505 mol) as white crystals in 85% isolated yield.

**^1^H NMR** (500 MHz, Acetone-*d*_6_) δ 8.06 (d, *J* = 9.4 Hz, 1H), 7.68 (d, *J* = 15.7 Hz, 1H), 7.63 (dd, *J* = 7.8, 1.8 Hz, 2H), 7.47 – 7.37 (m, 3H), 6.94 (d, *J* = 15.7 Hz, 1H), 6.78 – 6.69 (m, 1H), 6.18 – 6.08 (m, 1H) ppm; **^13^C-NMR:** (125 MHz, (CD_3_)_2_CO): 165.1, 141,7, 135.0, 129.8, 128.9, 127.8, 120.7, 102.2, 81.1 ppm. *Analytical data matches that reported in the literature.^2^*

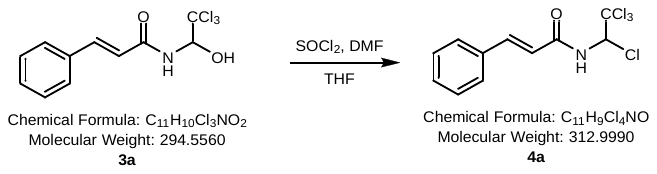

The reaction was carried out according to general procedure D using **3a** (148.72 g, 0.505 mol, 1 equiv.), SOCl_2_ (110 mL, 1.515 mol, 3 equiv.), DMF (1.96 mL, 25.25 mmol, 0.05 equiv.), and THF (505 mL, 1 M) to afford **4a** (150.16 g, 0.480 mol) as a light yellow powder in 95% isolated yield.

**^1^H-NMR**: (500 MHz, (CD_3_)_2_CO): 8.78 (d, *J=* 10.8 Hz, 1H), 7.75 (d, *J=* 15.4 Hz, 1H), 7.64 (m, 2H), 7.43 (m, 3H), 6.92 (d, *J=* 15.4 Hz, 1H), 6.82 (d, *J=* 10.8 Hz, 1H) ppm; ^13^**C-NMR:** (125 MHz, (CD_3_)_2_CO): 164.8, 143.5, 134.6, 130.3, 129.0, 128.1, 119.2, 99.6, 75.0 ppm. *Analytical data matches that reported in the literature.^2^*

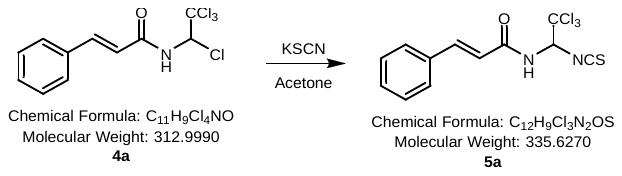

The reaction was carried out according to general procedure E using **4a** (150.16 g, 0.48 mol, 1 equiv.), potassium thiocyanate (46.65 g, 0.48 mol), and acetone (480 mL, 1 M) to afford **5a** (153.05 g, 0.456 mol) as a yellow solid in 95% isolated yield.

**^1^H-NMR**: (500 MHz, (CD_3_)_2_CO): 8.80 (d, *J=* 9 Hz, 1H), 7.76 (d, *J=* 16.1 Hz, 1H), 7.65 (m, 2H), 7.45 (m, 3H), 6.91 (d, *J=* 16.6 Hz, 1H), 6.61 (d, *J=* 9 Hz, 1H) ppm; **^13^C-NMR:** (125 MHz, (CD_3_)_2_CO): 165.2, 143.4, 142.1, 134.6, 130.3, 129.0, 128.1, 119.2, 99.3, 73.0 ppm. *Analytical data matches that reported in the literature.^2^*

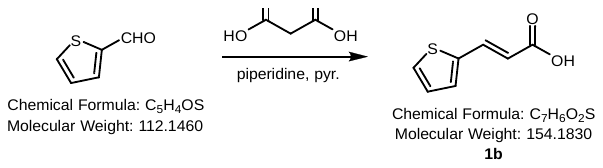

The reaction was carried out according to general procedure A using 2-thiophenecarboxaldehyde (4.67 mL, 50.0 mmol, 1 equiv.), malonic acid (10.41 g, 100.0 mmol, 2 equiv.), piperidine (0.49 mL, 5 mmol, 0.1 equiv.), and pyridine (100 mL, 0.5 M). Compound **1b** (6.47 g, 42.0 mmol) was obtained as a beige solid in 84% isolated yield.

**^1^H-NMR**: (500 MHz, CDCl_3_): δ 7.91 (d, *J=*15.6 Hz, 1H), 7.45 (d, *J=* 5.0 Hz, 1H), 7.33 (d, *J=* 3.5 Hz, 1H), 7.11 (dd, *J=* 5.0, 3.6 Hz, 1H), 6.27 (d, *J=* 15.6 Hz, 1H) ppm. *Analytical data matches that reported in the literature.^2^*

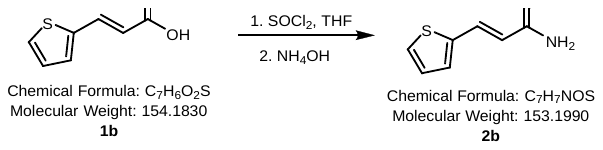

The reaction was carried out according to general procedure B using **1b** (4.63 g, 30.0 mmol, 1equiv.), SOCl_2_ (6.53 mL, 90.0 mmol, 3 equiv.), DMF (116 μL, 1.5 mmol, 0.05 equiv.), THF (100 mL, 0.3 M), and then NH_4_OH (53 mL, 5 equiv.) to afford **2b** (4.04 g, 26.4 mmol) in 88% isolated yield.

**^1^H-NMR**: (300 MHz, (CD_3_)_2_CO): δ 7.66 (d, *J=* 15.6 Hz, 1H), 7.51 (d, *J=* 5.2 Hz, 1H), 7.33 (d, *J=* 3.7 Hz, 1H), 7.10 (dd, *J=* 5.1, 3.6 Hz, 1H), 6.97 (bs, 1H), 6.47 (d, *J=* 15.1 Hz, 1H), 6.38 (bs, 1H) ppm. *Analytical data matches that reported in the literature.^2^*

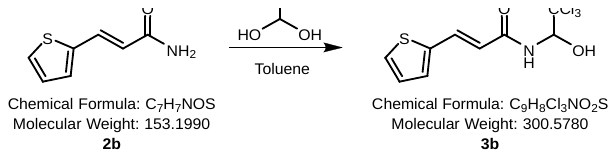

The reaction was carried out according to general procedure C using **2b** (3.06 g, 20 mmol, 1 equiv.), chloral hydrate (6.62 g, 40 mmol, 2 equiv.), and Toluene (40 mL, 0.5 M) to afford **3b** (5.53 g, 18.4 mmol) in 92% isolated yield.

**^1^H-NMR**: (300 MHz, (CD_3_)_2_CO): δ 8.08 (d, *J=* 9.1 Hz, 1H), 7.79 (d, *J=* 15.6 Hz, 1H), 7.56 (d, *J=* 5.2 Hz, 1H), 7.39 (d, *J=* 3.5 Hz, 1H), 7.12 (dd, *J=* 5.0, 3.7 Hz, 1H), 6.70 (d, *J=* 6.3 Hz, 1H), 6.67 (d, *J=* 15.6 Hz, 1H), 6.10 (dd, *J=* 9.3, 5.3 Hz, 1H) ppm. *Analytical data matches that reported in the literature.^2^*

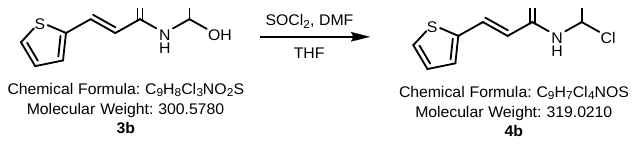

The reaction was carried out according to general procedure D using **3b** (4.51 g, 15 mmol, 1 equiv.), SOCl_2_ (3.26 mL, 45 mmol, 3 equiv.), DMF (58 μL, 0.75 mmol, 0.05 equiv.), and THF (50 mL, 0.3 M) to afford **4b** (4.54 g, 14.25 mmol) in 95% isolated yield.

**^1^H-NMR**: (300 MHz, (CD_3_)_2_CO): δ 8.77 (d, *J=* 10.5 Hz, 1H), 7.89 (d, *J=* 15.0 Hz, 1H), 7.61 (d, *J=* 5.2 Hz, 1H), 7.46 (d, *J=* 3.5 Hz, 1H), 7.14 (dd, *J=* 5.2, 3.7 Hz, 1H), 6.80 (d, *J=* 10.7 Hz, 1H), 6.65 (d, *J=* 15.2 Hz, 1H) ppm. *Analytical data matches that reported in the literature.^2^*

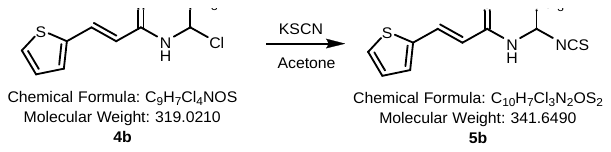

The reaction was carried out according general procedure E using **4b** (3.19 g, 10 mmol, 1 equiv.), KSCN (972 mg, 10 mmol, 1 equiv.), and acetone (20 mL, 0.5 M) to afford **5b** (3.25 g, 9.5 mmol) in a 95% isolated yield.

**^1^H-NMR**: (500 MHz, (CD_3_)_2_CO): δ 8.77 (d, *J=* 9.2 Hz, 1H), 7.89 (d, *J=* 14.5 Hz, 1H), 7.62 (d, *J=* 5.1 Hz, 1H), 7.47 (d, *J=* 3.4 Hz, 1H), 7.16 (dd, *J=* 5.1, 3.8 Hz, 1H), 6.65 (d, *J=* 15.4 Hz, 1H), 6.57 (d, *J=* 9.6 Hz, 1H) ppm. *Analytical data matches that reported in the literature.^2^*

#### b) Synthesis of sal003 Analogs in Figure 1

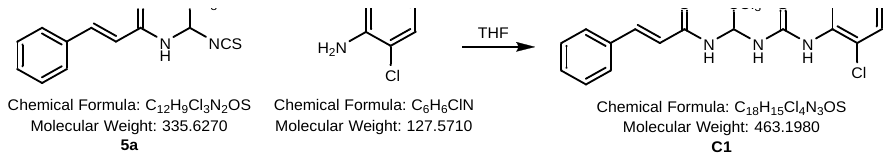

The reaction was carried out according to general procedure F using **5a** (1.58 g, 4.0 mmol, 1 equiv.), 2-chloroaniline (410 μL, 4.0 mmol, 1 equiv.), and THF (20 mL, 0.2 M) to afford **C1** (1.72 g, 3.7 mmol) as a white powder in 93% isolated yield.

**^1^H-NMR** (400 MHz, DMSO-*d*_6_) δ 9.97 (s, 1H), 9.08 (d, *J* = 8.8 Hz, 1H), 8.60 (d, *J* = 9.5 Hz, 1H), 7.74 (dd, *J* = 8.1, 1.6 Hz, 1H), 7.67 – 7.38 (m, 8H), 7.35 (td, *J* = 7.7, 1.5 Hz, 1H), 7.26 (td, *J* = 7.7, 1.7 Hz, 1H), 6.83 (d, *J* = 15.8 Hz, 1H) ppm; **^13^C NMR** (101 MHz, DMSO-*d*_6_) δ 182.8, 164.8, 141.5, 136.3, 135.0, 130.4, 129.9, 129.9, 129.5, 129.3, 128.2, 128.0, 127.5, 121.3, 102.0, 70.5 ppm. *Analytical data matches that reported in the literature.^2^*

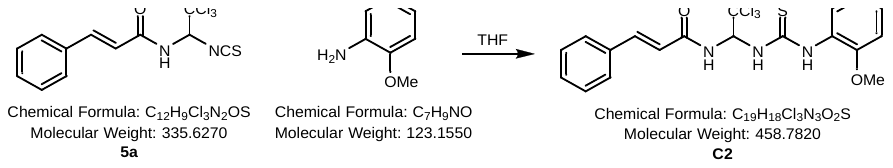

The reaction was carried out according to general procedure F using **5a** (1.58 g, 4.0 mmol, 1 equiv.), 2-methoxyaniline (451 μL, 4.0 mmol, 1 equiv.), and THF (20 mL, 0.2 M) to afford **C2** (1.65 g, 3.6 mmol) as a white powder in 90% isolated yield.

**^1^H-NMR** (400 MHz, DMSO-*d*_6_) δ 9.82 (s, 1H), 9.02 (d, *J* = 8.7 Hz, 1H), 8.47 (bs, 1H), 8.00 – 7.83 (m, 1H), 7.66 – 7.38 (m, 7H), 7.22 – 7.12 (m, 1H), 7.07 (dd, *J* = 8.3, 1.4 Hz, 1H), 6.94 (td, *J* = 7.6, 1.4 Hz, 1H), 6.82 (d, *J* = 15.8 Hz, 1H), 3.84 (s, 3H) ppm; **^13^C NMR** (101 MHz, DMSO-*d*_6_) δ 181.6, 164.7, 152.2, 141.4, 135.0, 130.3, 129.5, 128.2, 127.6, 126.6, 126.4, 121.4, 120.2, 111.9, 102.1, 70.3, 56.1 ppm. *Analytical data matches that reported in the literature.^2^*

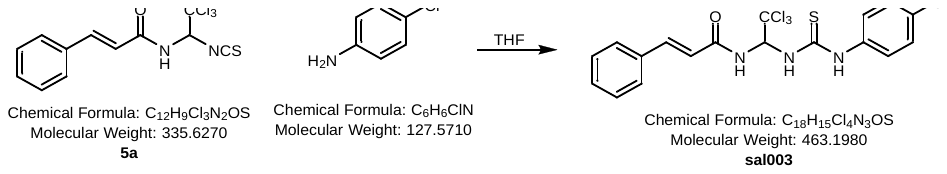

The reaction was carried out according to general procedure F using **5a** (1.58 g, 4.0 mmol, 1 equiv.), 4-chloroaniline (510 mg, 4.0 mmol, 1 equiv.), and THF (20 mL, 0.2 M) to afford **sal003** (1.67 g, 3.6 mmol) as a white powder in 90% isolated yield.

**^1^H-NMR** (400 MHz, DMSO-*d*_6_) δ 10.38 (s, 1H), 8.99 (d, *J* = 8.7 Hz, 1H), 8.27 (d, *J* = 9.5 Hz, 1H), 7.68 – 7.53 (m, 5H), 7.48 – 7.38 (m, 6H), 6.79 (d, *J* = 15.8 Hz, 1H) ppm; ^13^**C-NMR:** (125 MHz, DMSO-*d*_6_) δ 181.2, 164.8, 141.6, 138.3, 135.0, 130.4, 129.5, 129.1, 129.0, 128.3, 125.2, 121.3, 101.9, 70.1 ppm. *Analytical data matches that reported in the literature.^2^*

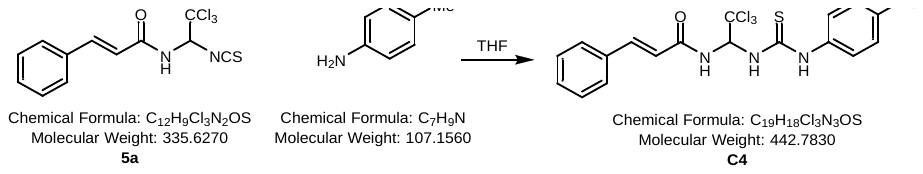

The reaction was carried out according to general procedure F using **5a** (1.58 g, 4.0 mmol, 1 equiv.), 4-methylaniline (429 mg, 4.0 mmol, 1 equiv.), and THF (20 mL, 0.2 M) to afford **C4** (1.56 g, 3.5 mmol) in 88% isolated yield.

**^1^H-NMR** (400 MHz, DMSO-*d*_6_) δ 10.25 (s, 1H), 8.97 (d, *J* = 8.6 Hz, 1H), 8.14 – 7.90 (m, 1H), 7.66 – 7.53 (m, 3H), 7.49 – 7.35 (m, 6H), 7.18 (d, *J* = 8.2 Hz, 2H), 6.76 (d, *J* = 15.8 Hz, 1H), 2.29 (s, 3H) ppm; **^13^C NMR** (101 MHz, DMSO-*d*_6_) δ 181.1, 164.7, 141.5, 136.5, 135.0, 134.8, 130.4, 129.7, 129.5, 128.2, 124.0, 121.3, 102.2, 70.2, 21.0 ppm. *Analytical data matches that reported in the literature.^2^*

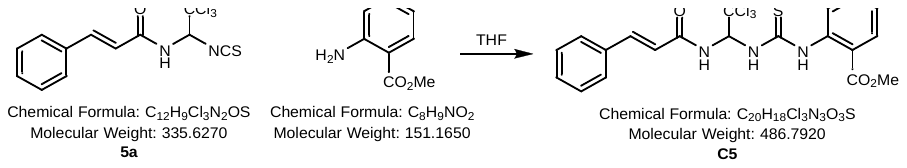

The reaction was carried out according to general procedure F using **5a** (1.58 g, 4.0 mmol, 1 equiv.), methyl 2-aminobenzoate (518 μL, 4.0 mmol, 1 equiv.), and THF (20 mL, 0.2 M) to afford **C5** (1.79 g, 3.7 mmol) in 92% isolated yield.

**^1^H-NMR** (400 MHz, DMSO-*d*_6_) δ 10.33 (s, 1H), 9.02 (d, *J* = 8.9 Hz, 1H), 8.92 (d, *J* = 9.4 Hz, 1H), 7.83 (ddd, *J* = 23.9, 8.0, 1.4 Hz, 2H), 7.67 – 7.37 (m, 8H), 7.31 (td, *J* = 7.6, 1.2 Hz, 1H), 6.88 (d, *J* = 15.8 Hz, 1H), 3.81 (s, 3H) ppm; **^13^C NMR** (101 MHz, DMSO-*d*_6_) δ 182.8, 166.7, 164.8, 141.5, 139.6, 135.1, 132.8, 130.6, 130.4, 129.5, 128.3, 128.2, 125.8, 124.5, 121.4, 101.9, 70.5, 52.8 ppm. *Analytical data matches that reported in the literature.^2^*

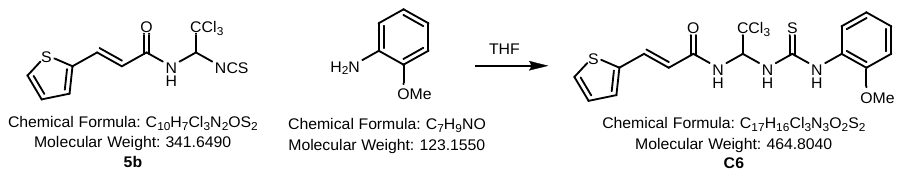

The reaction was carried out according to general procedure F using **5b** (1.37 g, 4 mmol, 1 equiv.), 2-methoxylaniline (451 μL, 4.0 mmol, 1 equiv.), and THF (20 mL, 0.2 M) to afford **C6** (1.67 g, 3.6 mmol) as a white powder in 91% isolated yield.

**^1^H-NMR** (500 MHz, DMSO-*d*_6_) δ 9.81 (s, 1H), 9.00 (d, *J* = 8.8 Hz, 1H), 8.49 (s, 1H), 7.90 (d, *J* = 7.7 Hz, 1H), 7.70 (d, *J* = 15.5 Hz, 1H), 7.64 (d, *J* = 5.1 Hz, 1H), 7.49 – 7.41 (m, 2H), 7.18 – 7.10 (m, 2H), 7.05 (dd, *J* = 8.4, 1.3 Hz, 1H), 6.92 (td, *J* = 7.7, 1.3 Hz, 1H), 6.56 (d, *J* = 15.5 Hz, 1H), 3.82 (s, 3H) ppm; **^13^C NMR** (126 MHz, DMSO-*d*_6_) δ 181.6, 164.5, 152.1, 140.0, 134.4, 131.9, 129.1, 128.9, 127.6, 126.6, 126.4, 120.2, 119.9, 111.9, 102.1, 70.2, 56.1 ppm. *Analytical data matches that reported in the literature.^2^*

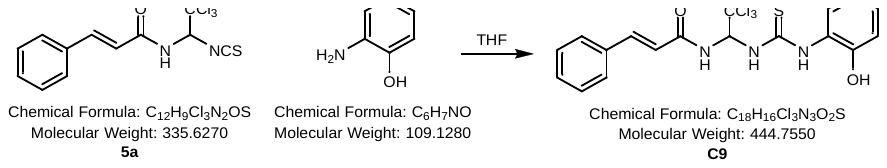

The reaction was carried out according to general procedure F using **5a** (1.58 g, 4.0 mmol, 1 equiv.), 2-aminophenol (437 mg, 4.0 mmol, 1 equiv.), and THF (20 mL, 0.2 M) to afford **C9** (1.51 g, 3.4 mmol) in 85% isolated yield.

**R_f_ =** (EtOAc/hexanes 1:1): 0.24; **IR** (neat) v = 3287.4, 3210.6, 3083.0, 3026.1, 2955.5, 1494.7, 1452.0, 1342.6, 1206.4, 1136.0, 742.0 cm^-1^; **^1^H NMR** (400 MHz, DMSO-*d*_6_) δ 9.93 (s, 1H), 9.78 (s, 1H), 9.02 (d, *J* = 8.8 Hz, 1H), 8.55 (bs, 1H), 8.02 – 7.81 (m, 1H), 7.66 – 7.35 (m, 7H), 7.08 – 6.72 (m, 4H) ppm; **^13^C NMR** (101 MHz, DMSO-*d*_6_) δ 181.4, 164.7, 150.1, 141.4, 135.1, 130.3, 129.5, 128.2, 126.7, 126.0, 121.4, 118.8, 115.8, 102.1, 70.3 ppm; **HRMS:** Calcd. for C_18_H_16_Cl_3_N_3_NaO_2_S: [M+Na]^+^: 465.9921 m/z, found 465.9920 m/z.

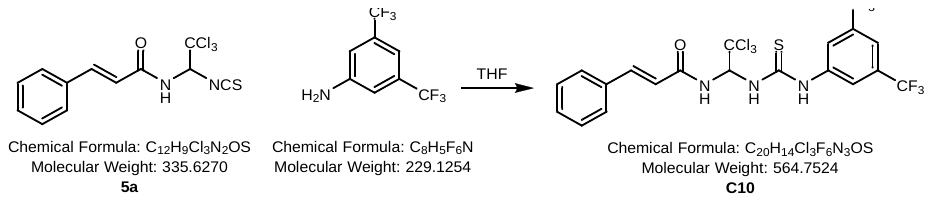

The reaction was carried out according to general procedure F using **5a** (1.58 g, 4.0 mmol, 1 equiv.), 3,5-Bis(trifluoromethyl)aniline (625 μL, 4.0 mmol, 1 equiv.), and THF (20 mL, 0.2 M) to afford **C10** (2.10 g, 3.7 mmol) in 93% isolated yield.

**R_f_ =** (EtOAc/hexanes 1:1): 0.51; **IR** (neat) v = 3232.1, 3070.2, 2998.7, 2945.5, 1656.0, 1621.6, 1503.4, 1376.5, 1274.3, 1174.9, 1131.1, 954.8, 896.3, 837.2, 811.3, 767.7, 681.1, 490.1 cm^-1^; **^1^H-NMR** (400 MHz, DMSO-*d*_6_) δ 10.79 (s, 1H), 9.03 (d, *J* = 8.8 Hz, 1H), 8.62 (d, *J* = 9.4 Hz, 1H), 8.33 (s, 2H), 7.84 (s, 1H), 7.68 – 7.53 (m, 3H), 7.43 (q, *J* = 8.1, 7.3 Hz, 4H), 6.84 (d, *J* = 15.8 Hz, 1H) ppm; **^13^C NMR** (101 MHz, DMSO-*d*_6_) δ 181.6, 164.9, 141.7, 141.6, 135.0, 131.3, 131.0, 130.7, 130.4, 129.5, 128.3, 127.6, 124.9, 123.1, 122.2, 121.2, 119.5, 117.9, 101.7, 70.0 ppm; **HRMS:** Calcd. for C_20_H_14_Cl_3_F_6_N_3_NaOS: [M+Na]^+^: 585.9720 m/z, found 585.9712 m/z.

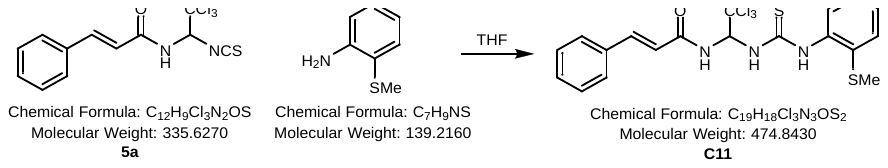

The reaction was carried out according to general procedure F using **5a** (1.58 g, 4.0 mmol, 1 equiv.), 2-(methylthio)aniline (501 μL, 4.0 mmol, 1 equiv.), and THF (20 mL, 0.2 M) to afford **C11** (1.80 g, 3.8 mmol) in 95% isolated yield.

**R_f_ =** (EtOAc/hexanes 1:1): 0.36; **IR** (neat) v = 3311.1, 3250.4, 3033.9, 1654.4, 1608.8, 1492.0, 1274.5, 1090.5, 1072.1, 992.0, 887.1, 766.7, 711.5, 686.8, 602.5, 513.4, 480.3 cm^-1^; **^1^H NMR** (400 MHz, DMSO-*d*_6_) δ 9.76 (s, 1H), 9.07 (d, *J* = 8.8 Hz, 1H), 8.35 (s, 1H), 7.66 – 7.53 (m, 3H), 7.52 – 7.38 (m, 5H), 7.35 – 7.24 (m, 2H), 7.18 (td, *J* = 7.5, 1.6 Hz, 1H), 6.83 (d, *J* = 15.8 Hz, 1H), 2.42 (s, 3H) ppm; **^13^C NMR** (101 MHz, DMSO-*d*­_6_) δ 183.0, 164.7, 141.5, 136.4, 136.0, 135.0, 130.4, 129.5, 129.3, 128.2, 127.7, 126.6, 125.3, 121.4, 102.2, 70.5, 15.1 ppm; **HRMS:** Calcd. for C_19_H_18_Cl_3_N_3_NaOS_2_: [M+Na]^+^: 495.9849 m/z, found 495.9862 m/z.

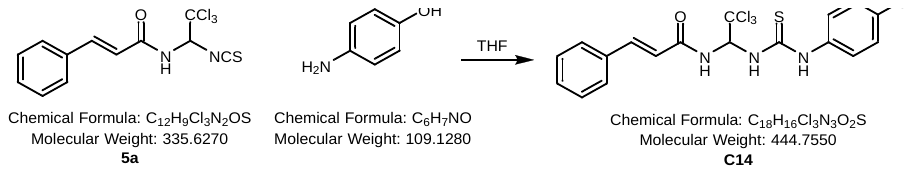

The reaction was carried out according to general procedure F using **5a** (1.58 g, 4.0 mmol, 1 equiv.), 4-aminophenol (437 mg, 4.0 mmol, 1 equiv.), and THF (20 mL, 0.2 M) to afford **C14** (1.51 g, 3.4 mmol) in 85% isolated yield.

**R_f_ =** (EtOAc/hexanes 1:1): 0.16; **IR** (neat) v = 3197.9, 3083.5, 3025.7, 2957.1, 1654.9, 1617.1, 1500.8, 1341.2, 1203.5, 1030.9, 887.7, 792.4, 764.0, 682.6, 554.1 cm^-1^; **^1^H NMR** (500 MHz, DMSO-*d*_6_) δ 10.06 (s, 1H), 9.49 (s, 1H), 8.95 (d, *J* = 8.7 Hz, 1H), 8.19 – 7.49 (m, 4H), 7.49 – 7.36 (m, 4H), 7.20 (d, *J* = 8.0 Hz, 2H), 6.75 (dd, *J* = 22.7, 11.9 Hz, 3H) ppm; **^13^C NMR** (101 MHz, DMSO-*d*_6_) δ 181.3, 164.6, 155.8, 141.5, 135.0, 134.9, 130.4, 129.5, 128.3, 126.4, 121.3, 115.8, 102.3, 70.3 ppm; **HRMS:** Calcd. for C_18_H_16_Cl_3_N_3_NaO_2_S: [M+Na]^+^: 465.9921 m/z, found 465.9911 m/z.

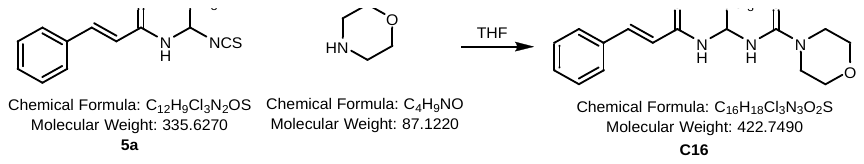

The reaction was carried out according to general procedure F using **5a** (1.58 g, 4.0 mmol, 1 equiv.), morpholine (345 μL, 4.0 mmol, 1 equiv.), and THF (20 mL, 0.2 M) to afford **C16** (1.62 g, 3.84 mmol) in 96% isolated yield.

**^1^H NMR** (500 MHz, DMSO-*d*_6_) δ 8.36 (d, *J* = 8.9 Hz, 1H), 7.89 (d, *J* = 8.7 Hz, 1H), 7.68 – 7.63 (m, 2H), 7.61 (t, *J* = 8.8 Hz, 1H), 7.56 (d, *J* = 15.8 Hz, 1H), 7.47 – 7.39 (m, 3H), 6.80 (d, *J* = 15.7 Hz, 1H), 3.93 – 3.82 (m, 2H), 3.83 – 3.73 (m, 2H), 3.64 (pt, *J* = 6.5, 3.6 Hz, 4H) ppm; **^13^C NMR** (126 MHz, DMSO-*d*_6_) δ 182.7, 164.6, 141.7, 134.9, 130.4, 129.4, 128.4, 121.4, 102.7, 71.6, 66.1, 49.0 ppm. *Analytical data matches that reported in the literature.^2^*

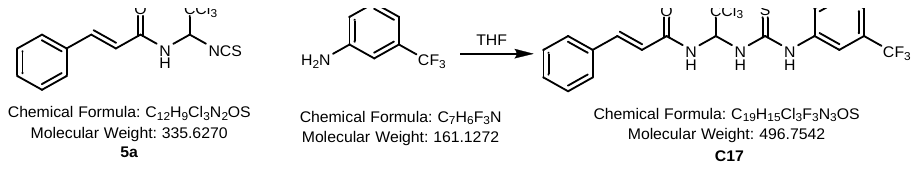

The reaction was carried out according to general procedure F using **5a** (1.58 g, 4.0 mmol, 1 equiv.), 3-(trifluoromethyl)aniline (494 μL, 4.0 mmol, 1 equiv.), and THF (20 mL, 0.2 M) to afford **C17** (1.82 g, 3.76 mmol) in 94% isolated yield.

**R_f_ =** (EtOAc/hexanes 1:1): 0.40; **IR** (neat) v = 3195.7, 3092.7, 1654.3, 1617.5, 1505.2, 1328.6, 1164.4, 1127.9, 1040.4, 967.4, 931.5, 831.5, 770.5, 716.1, 619.4, 560.3 cm^-1‑^; **^1^H NMR** (500 MHz, DMSO-*d*_6_) δ 10.57 (s, 1H), 9.01 (d, *J* = 8.8 Hz, 1H), 8.42 (d, *J* = 9.5 Hz, 1H), 8.14 (d, *J* = 1.9 Hz, 1H), 7.77 (dd, *J* = 8.1, 2.0 Hz, 1H), 7.64 – 7.38 (m, 9H), 6.81 (d, *J* = 15.8 Hz, 1H) ppm; **^13^C NMR** (126 MHz, DMSO-*d*_6_) δ 181.0, 164.8, 141.6, 140.3, 135.0, 130.4, 130.3, 129.8, 129.5, 129.5, 128.3, 127.0, 125.5, 123.4, 121.5, 121.2, 119.5, 101.9, 70.0 ppm; **HRMS:** Calcd. for C_19_H_15_Cl_3_F_3_N_3_NaOS: [M+Na]^+^: 517.9846 m/z, found 517.9856 m/z.

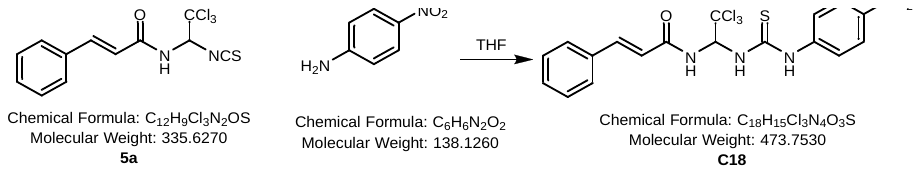

The reaction was carried out according to general procedure F using **5a** (1.58 g, 4.0 mmol, 1 equiv.), 4-nitroaniline (552 mg, 4.0 mmol, 1 equiv.), and THF (20 mL, 0.2 M) to afford **C18** (1.68 g, 3.52 mmol) in 88% isolated yield.

**R_f_ =** (EtOAc/hexanes 1:1): 0.26; **IR** (neat) v = 3283.5, 3196.7, 3080.9, 3025.9, 2938.4, 1661.1, 1629.3, 1492.5, 1330.7, 1202.7, 1111.6, 969.3, 892.4, 802.0, 765.1, 703.5, 545.9 cm^-1^; **^1^H NMR** (400 MHz, DMSO-*d*_6_) δ 10.86 (s, 1H), 9.07 (d, *J =* 9.4 Hz, 1H), 8.65 (d, *J* = 9.4 Hz, 1H), 8.23 (d, *J* = 9.2 Hz, 2H), 8.01 (d, *J* = 9.2 Hz, 2H), 7.67 – 7.53 (m, 3H), 7.53 – 7.34 (m, 4H), 6.82 (d, *J* = 15.8 Hz, 1H) ppm; **^13^C NMR** (101 MHz, DMSO-*d*_6_) δ 180.8, 164.9, 145.9, 143.2, 141.7, 135.0, 130.4, 129.5, 128.3, 125.0, 121.8, 121.2, 101.6, 69.9 ppm; **HRMS:** Calcd. for C_18_H_15_Cl_3_N_4_NaO_3_S: [M+Na]^+^: 494.9823 m/z, found 494.9804 m/z.

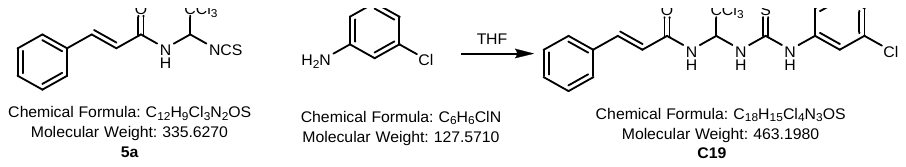

The reaction was carried out according to general procedure F using **5a** (1.58 g, 4.0 mmol, 1 equiv.), 3-chloroaniline (423 μL, 4.0 mmol, 1 equiv.), and THF (20 mL, 0.2 M) to afford **C19** (1.77 g, 3.84 mmol) in 96% isolated yield.

**R_f_ =** (EtOAc/hexanes 1:1): 0.45; **IR** (neat) v = 3260.8, 3190.9, 3089.4, 3057.0, 2993.5, 1654.1, 1618.3, 1502.1, 1337.1, 1206.9, 1131.2, 1103.7, 1040.0, 966.7, 827.5, 806.9, 764.7, 706.6, 559.5 cm^-1^; **^1^H NMR** (400 MHz, DMSO-*d*_6_) δ 10.44 (s, 1H), 9.00 (d, *J* = 8.7 Hz, 1H), 8.35 (d, *J* = 9.4 Hz, 1H), 7.88 (s, 1H), 7.66 – 7.54 (m, 3H), 7.42 (dt, *J* = 14.4, 7.9 Hz, 6H), 7.22 (d, *J* = 6.9 Hz, 1H), 6.80 (d, *J* = 15.8 Hz, 1H) ppm; **^13^C NMR** (101 MHz, DMSO-*d*_6_) δ 181.2, 164.8, 141.6, 140.9, 135.0, 133.2, 130.8, 130.4, 129.5, 128.3, 125.0, 122.8, 121.8, 121.3, 101.9, 70.1 ppm; **HRMS:** Calcd. for C_18_H_15_Cl_4_N_3_NaOS: [M+Na]^+^: 483.9582 m/z, found 483.9576 m/z.

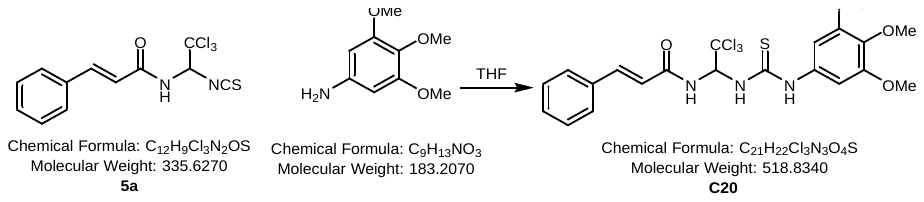

The reaction was carried out according to general procedure F using **5a** (1.58 g, 4.0 mmol, 1 equiv.), 3,4,5-trimethoxyaniline (733 mg, 4.0 mmol, 1 equiv.), and THF (20 mL, 0.2 M) to afford **C20** (1.86 g, 3.6 mmol) in 90% isolated yield.

**R_f_ =** (EtOAc/hexanes 1:1): 0.23; **IR** (neat) v = 3220.7, 3067.5, 2942.2, 2830.4, 1502.5, 1334.3, 1229.1, 1167.4, 1129.9, 1097.9, 1043.0, 1007.6, 840.7, 811.5, 718.1, 690.7 cm^-1^; **^1^H NMR** (400 MHz, DMSO-*d*_6_) δ 10.31 (s, 1H), 8.91 (d, *J* = 8.7 Hz, 1H), 8.04 (d, *J* = 8.9 Hz, 1H), 7.66 – 7.59 (m, 2H), 7.56 (d, *J* = 15.8 Hz, 1H), 7.49 – 7.37 (m, 4H), 6.83 (s, 2H), 6.74 (d, *J* = 15.8 Hz, 1H), 3.77 (s, 6H), 3.66 (s, 3H) ppm; **^13^C NMR** (101 MHz, DMSO-*d*_6_) δ 180.7, 164.6, 153.2, 141.5, 135.3, 134.9, 134.6, 130.4, 129.5, 128.3, 121.3, 102.2, 101.6, 70.2, 60.5, 56.3 ppm; **HRMS:** Calcd. for C_21_H_22_Cl_3_N_3_NaO_4_S: [M+Na]^+^: 540.0289 m/z, found 540.0286 m/z.

#### c.) Synthesis of C7a

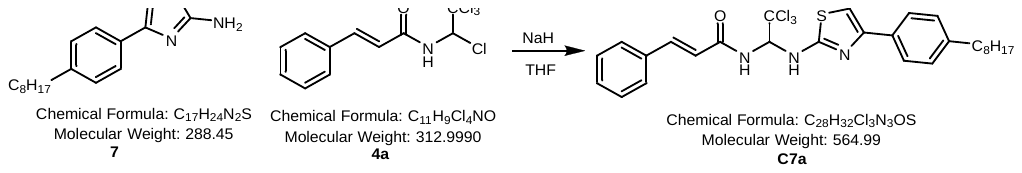

In a flame-dried 25 mL round bottom flask equipped with a Teflon-coated stir bar, **7** (150 mg, 0.52 mmol, 1 eq.) was dissolved in anhydrous THF (4 mL) and cooled in an ice-bath. NaH (60% dispersion in mineral oil, 20 mg, 0.52 mmol, 1 eq.) was added slowly and the mixture was stirred for 15 minutes. Chloral amide **4a** (163 mg, 0.52 mmol, 1 eq.) was dissolved in anhydrous THF (3 mL) and added rapidly to the reaction mixture, which was then stirred for 1 hour at rt. The solvent was removed *in vacuo* and the crude mixture was purified using silica gel chromatography (9:1 DCM/hexanes) to yield **C7a** (132 mg, 0.234 mmol) as a yellow solid in 45%.

**^1^H NMR** (500 MHz, DMSO-*d*_6_) δ 8.98 (d, *J* = 9.1 Hz, 1H), 8.63 (d, *J* = 9.0 Hz, 1H), 7.75 (d, *J* = 8.3 Hz, 2H), 7.61 – 7.53 (m, 3H), 7.46 – 7.38 (m, 3H), 7.20 (d, *J* = 8.3 Hz, 2H), 7.13 (s, 1H), 7.00 – 6.91 (m, 2H), 2.58 (t, *J* = 7.6 Hz, 2H), 1.57 (t, *J* = 7.4 Hz, 2H), 1.34 – 1.18 (m, 10H), 0.90 – 0.81 (m, 3H) ppm.

#### d.) Synthesis of C7c

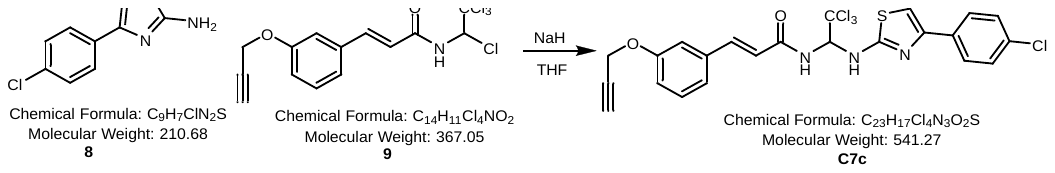

In a flame-dried 25 mL round bottom flask equipped with a Teflon-coated stir bar, **8** (100.5 mg, 0.48 mmol, 1 eq.) was dissolved in anhydrous THF (4 mL) and cooled in an ice-bath. NaH (60% dispersion in mineral oil, 18.3 mg, 0.48 mmol, 1 eq.) was added slowly and the mixture was stirred for 15 minutes. Chloral amide **9** (175 mg, 0.48 mmol, 1 eq.) was dissolved in anhydrous THF (3 mL) and added rapidly to the reaction mixture, which was then stirred for 1 hour at rt. The solvent was removed *in vacuo* and the crude mixture was purified using silica gel chromatography (9:1 DCM/hexanes) to yield **C7c** (91 mg, 0.168 mmol) as a yellow solid in 35%.

**^1^H NMR** (500 MHz, DMSO-*d*_6_) δ 8.90 (s, 1H), 8.68 (d, *J* = 9.0 Hz, 1H), 7.91 – 7.82 (m, 2H), 7.61 – 7.43 (m, 5H), 7.29 (s, 1H), 7.04 (d, *J* = 8.8 Hz, 2H), 6.97 (t, *J* = 9.0 Hz, 1H), 6.79 (d, *J* = 15.7 Hz, 1H), 4.85 (d, *J* = 2.4 Hz, 2H), 3.60 (t, *J* = 2.4 Hz, 1H) ppm.

#### e.) Synthesis of C8

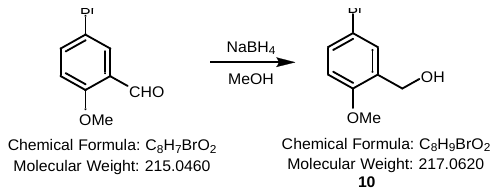

In a 250 mL round bottom flask equipped with a Teflon coated stir bar, 5-bromo-2-methoxybenzaldehyde (2.15g, 10 mmol, 1 eq.) was dissolved in MeOH (50 mL) and cooled in an ice bath. NaBH_4_ (95 mg, 2.5 mmol, 0.25 eq.) was added slowly and the mixture was stirred at rt for 1 hour. The reaction mixture was diluted with 1M HCl (10 mL) and extracted with DCM (3 x 50 mL). The organic layers were combined, washed with brine (50 mL), dried over MgSO_4_, filtered, and concentrated *in vacuo* to yield **10** (2.06 g, 9.5 mmol) as a white solid in 95% that was used without further purification.

**^1^H NMR** (300 MHz, CDCl_3_) δ 7.42 (d, *J* = 2.5 Hz, 1H), 7.37 (dd, *J* = 2.5 Hz, 8.7 Hz, 1H), 6.75 (d, *J* = 8.7 Hz, 1H), 4.64 (s, 2H), 3.84 (s, 3H), 1.87 (s, 2H) ppm; **^13^C NMR** (125 MHz, CDCl_3_) δ 188.5, 156.3, 138.3, 131.3, 112.9, 111.8, 61.1, 55.6 ppm. *Analytical data matches that reported in the literature.^3^*

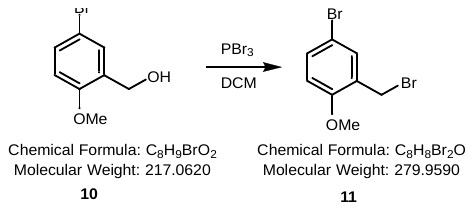

In a 250 mL round-bottom flask equipped with a Teflon coated stir-bar, **10** (2.06 g, 9.5 mmol, 1.0 eq) was dissolved in DCM (100 mL) and cooled to 0^˚^C in an ice bath. In a separate 50 mL round bottom flask, PBr_3_ (1.79 mL, 19 mmol, 2.0 eq) was dissolved in DCM (10 mL) and slowly added to the cooled solution, and then stirred at rt for 15 minutes. The reaction mixture was then concentrated *in vacuo* and the resulting slurry was carefully quenched with a cold, saturated NaHCO_3_ solution (100 mL). The reaction mixture was then extracted with DCM (3 x 30 mL), dried over MgSO_4_, filtered, and concentrated *in vacuo* to obtain the alkyl bromide **11** (2.34 g, 8.36 mmol) in 88% which was used without further purification.

**^1^H NMR** (300 MHz, CDCl_3_) δ 7.47 (d, *J* = 2.5 Hz, 1H), 7.41 (dd, *J* = 2.5 Hz, 8.7 Hz, 1H), 6.78 (d, *J =* 8.7 Hz, 1H), 4.50 (s, 2H), 3.90 (s, 3H) ppm; **^13^C NMR** (125 MHz, CDCl_3_) δ 156.5, 133.4, 132.7, 128.2, 112.7, 112.5, 55.9, 27.6 ppm. *Analytical data matches that reported in the literature.^4^*

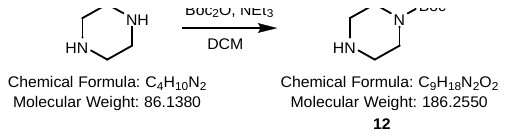

In a 500 mL round-bottom flask equipped with a Teflon coated stir-bar, piperazine (92.87 mmol, 2.0 eq) was dissolved in DCM (210 mL) and cooled to 0^˚^C in an ice-bath. Boc_2_O (46.43 mmol, 1.0 eq) was dissolved in DCM (20 mL), then added to reaction mixture dropwise and stirred for 1 hour. The reaction mixture was gravity-filtered, washed with cold DCM (2 x 30 mL), and concentrated *in vacuo*. Water (75 mL) was then added, and the resulting mixture was gravity filtered. The solution was saturated with K_2_CO_3_ before being extracted with EtOAc (3 x 30mL), dried over MgSO_4_, filtered, and concentrated *in vacuo* to yield the Boc-protected piperazine **12** (12.97 g, 69.65 mmol) in 75% as a white solid.

**^1^H NMR** (300 MHz, CDCl_3_) δ 3.38 (m, 4H), 2.79 (m, 4H), 1.80 (s, 1H), 1.45 (s, 9H).

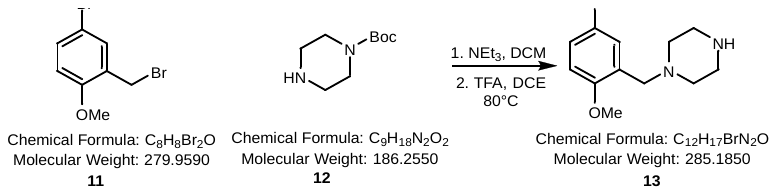

In a 250 mL round-bottom flask equipped with a Teflon coated stir-bar, **11** (9.07 mmol, 1.0 eq), **12** (9.07 mmol, 1.0 eq), and NEt_3_ (8.841 mmol, 1.0 eq) were dissolved in 1,2-dichloroethane (90 mL) and refluxed for 18 hours at 85^˚^C. The mixture was then washed with water (3 x 30 mL), dried over MgSO_4_, filtered, and concentrated *in vacuo*. The crude mixture was then dissolved in a 1,2-dichloroethane/TFA (1:1) mixture (20 mL) and heated at 80°C for 12 hours. After cooling to rt, the mixture was carefully washed with sat. NaHCO_3_ (50 mL) and brine (50 mL), dried over MgSO_4_, filtered and concentrated *in vacuo* to yield **13** (2.38 g, 8.34 mmol) in 92% as a white powder which was used without further purification.

**^1^H NMR** (300 MHz, DMSO-*d*_6_) δ 7.46-7.36 (m, 2H), 6.95 (d, *J* = 8.6 Hz, 1H), 3.75 (s, 3H), 3.48 (s, 2H), 3.30 (s, 4H), 2.98 (m, 4H), 1.36 (s, 1H) ppm.

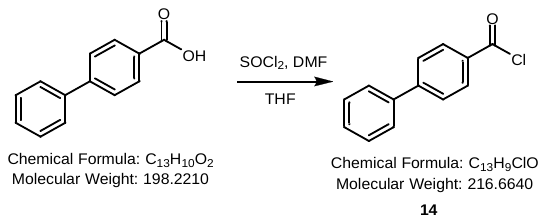

Was prepared according to the procedure for compound **4a**. Biphenyl-4-carboxylic acid (1.98 g, 10 mmol, 1 eq.) was dissolved in anhydrous THF (50 mL), followed by the addition of DMF and SOCl_2_ (2.2 ml, 30 mmol, 3 eq.). The mixture was heated at reflux for 2 hours, cooled to rt and the solvent removed *in vacuo* to yield the acid chloride **14** (2.06 g, 9.5 mmol) which was used in the next step without purification.

**^1^H NMR** (500 MHz, Acetone-*d*_6_) δ 8.27 – 8.23 (m, 2H), 7.99 – 7.93 (m, 2H), 7.83 – 7.79 (m, 2H), 7.58 – 7.54 (m, 2H), 7.52 – 7.48 (m, 1H) ppm.

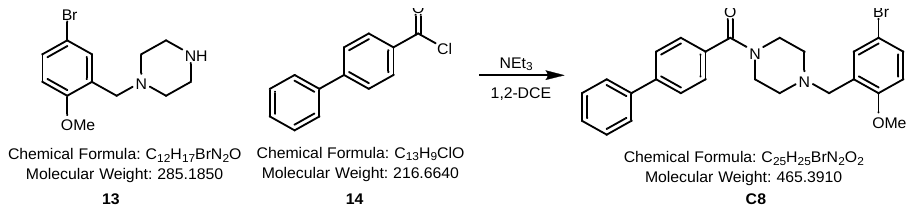

In a flame-dried 250 mL round bottom flask equipped with a Teflon coated stir bar, **13** (570 mg, 2 mmol, 1 eq.) was dissolved in anhydrous 1,2-dichloroethane (20 mL), NEt_3_ ­(0.51 mL, 4 mmol, 2 eq.) was added and the mixture was stirred at 60ºC for 30 minutes. The mixture was cooled to rt, **14** (433 mg, 2 mmol, 1 eq.) was added and the reaction was stirred and heated at reflux for 6 hours. After cooling to rt, the mixture was washed with 1M HCl (20 mL), dried with MgSO_4_, filtered and concentrated *in vacuo*. The crude mixture was purified using silica gel chromatography to yield **C8** (707 mg, 1.52 mmol) as a yellow solid in 76%.

**^1^H NMR** (500 MHz, Acetone-*d*_6_) δ 7.72 (dd, *J* = 12.6, 7.8 Hz, 4H), 7.60 – 7.46 (m, 5H), 7.44 – 7.36 (m, 2H), 6.96 (d, *J* = 8.7 Hz, 1H), 3.85 (s, 3H), 3.66 (bs, 2H), 3.58 (s, 2H), 2.52 (s, 4H), 1.30 (s, 2H) ppm; **^13^C NMR** (126 MHz, Acetone-*d*_6_) δ 169.0, 157.1, 142.0, 140.1, 135.5, 132.2, 130.6, 128.9, 128.9, 127.9, 127.7, 126.9, 126.7, 112.7, 112.2, 55.3, 55.0, 53.0 ppm. *Analytical data matches that reported in the literature.^5^*

#### f.) Synthesis of C13

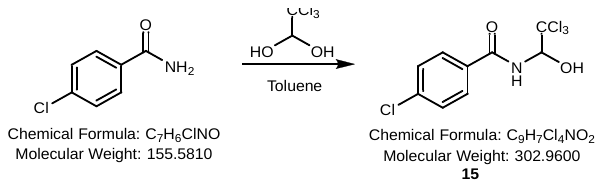

Compound **15** was prepared according to the procedure for **3a** using 4-chlorobenzamide (1.56 g, 10 mmol, 1.0 eq), chloral hydrate (3.31 g, 20 mmol, 2.0 eq), and toluene (100 mL) to obtain **15** (1.97 g, 6.5 mmol) as white crystals in 65%.

**^1^H NMR** (300 MHz, (CD_3_)_2_CO) δ 8.33 (d, *J* = 8.3 Hz, 1H), 7.98 (d, *J* = 8.6 Hz, 2H), 7.53 (d, *J* = 8.8 Hz, 2H), 6.85 (s, 1H), 6.21 (d, *J* = 9 Hz, 1H). **^13^C NMR** (125 MHz, (CD_3_)_2_CO) δ 165.8, 132.5, 129.5, 128.8, 128.6, 102.1, 81.7.

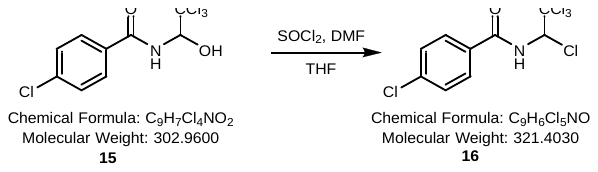

Was prepared according to the procedure for **4a** using **15** (1.42 g, 4.70 mmol, 1.0 eq), SOCl_2_ (1.02 mL, 14.11 mmol, 3.0 eq), DMF (catalytic amount, 1 mol%), and THF (50 mL) to obtain **16** (1.24 g, 3.85 mmol) as a yellow powder in 82%.

**^1^H NMR** (300 MHz, (CD_3_)_2_CO) δ 9.07 (d, *J* = 9.8 Hz, 1H), 7.98 (d, *J* = 8.8Hz, 2H), 7.57 (d, *J* = 8.8 Hz, 2H), 6.89 (d, *J* = 10.2 Hz, 1H). **^13^C NMR** (125 MHz, (CD_3_)_2_CO) δ 165.7, 138.3, 131.3, 129.9, 128.7, 99.7, 74.8.

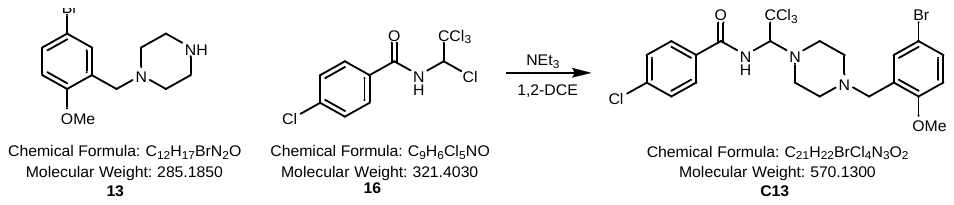

Was prepared according to the procedure for **C8** using **13** (200 mg, 0.70 mmol, 1.0 eq), **16** (225 mg, 0.70 mmol, 1.0 eq), NEt_3_ (89 μL, 0.70 mmol, 1.0 eq), and DCE (10 mL) to obtain The crude product was purified using a column of 1:1 EtOAc:Hexanes and concentrated *in vacuo*.

**Rf** = (EtOAc/hexanes 1:1) 0.6; **^1^H NMR** (500 MHz, (CD_3_)_2_CO) δ 8.23 (d, *J* = 9.5 Hz, 1H), 7.96 (d, *J* = 8.5 Hz, 2H), 7.54 (m, 3H), 7.37 (dd, *J* = 8.7, 2.5 Hz, 1H), 6.94 (d, *J* = 8.7 Hz, 1H), 5.63 (d, *J* = 9.6 Hz, 1H), 3.84 (s, 3H), 3.55 (s, 2H), 3.21 (s, 2H), 2.91 (s, 2H), 2.59 (s, 4H) ppm; **^13^C NMR** (125 MHz, (CD_3_)_2_CO) δ 166.8, 157.2, 137.4, 132.5, 130.8, 129.7, 128.5, 112.7, 112.2, 102.6, 79.7, 55.3, 54.7, 52.9, 29.7 ppm; **HRMS**: Calcd for C_21_H_23_BrCl_4_N_3_O_2_ (M+H)^+^: 567.97278 m/z, found 567.97223 m/z.

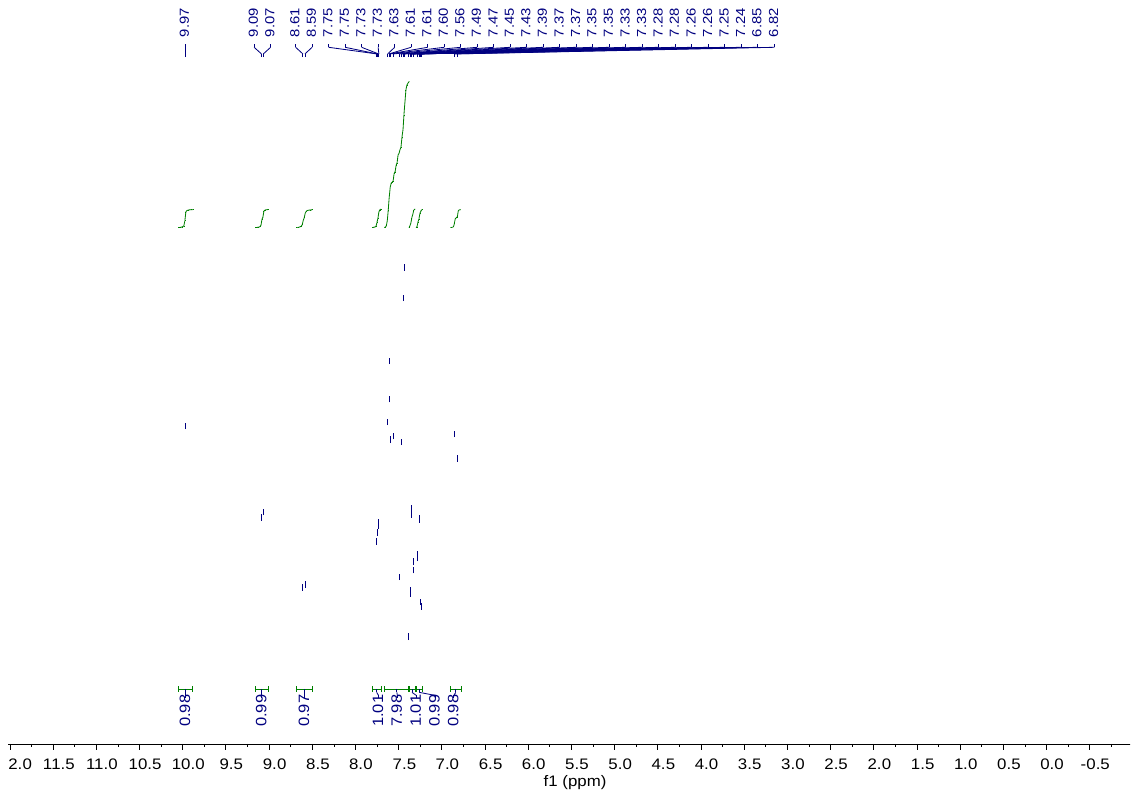

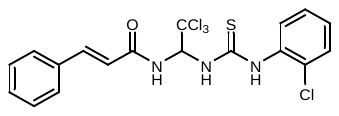
**C1**

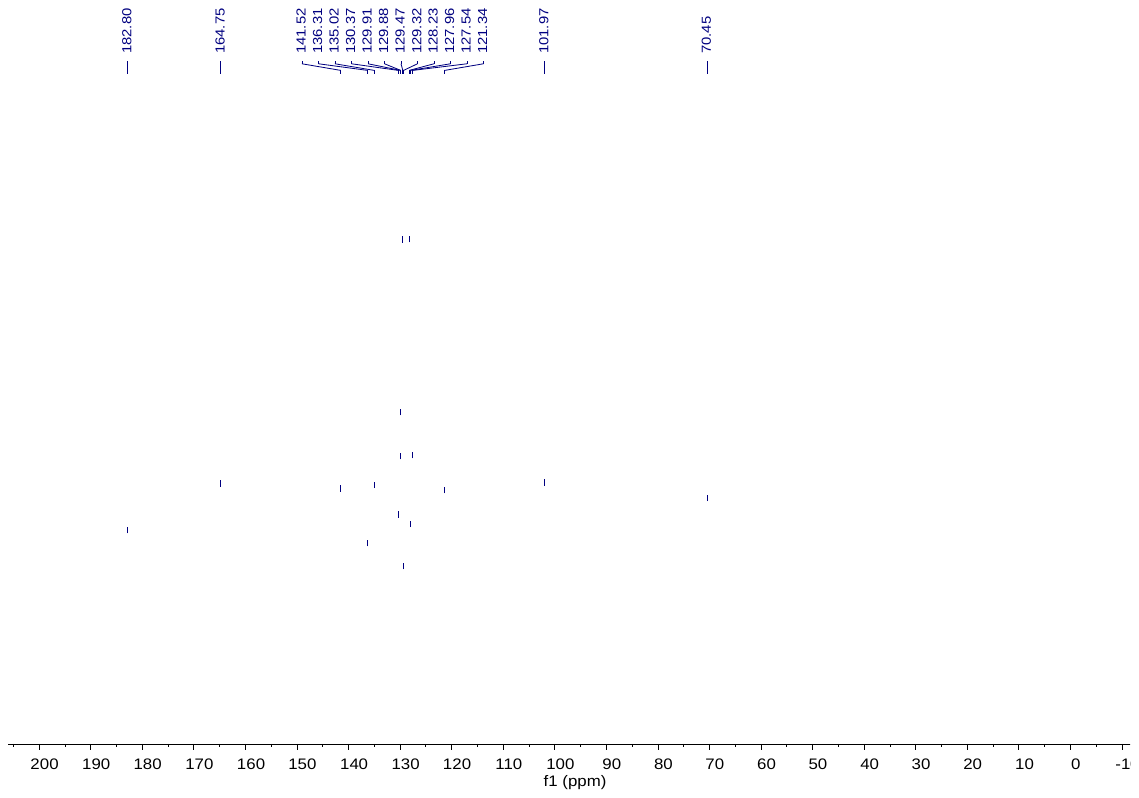
**C1**

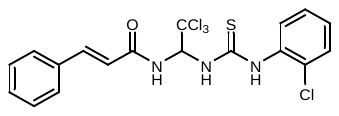

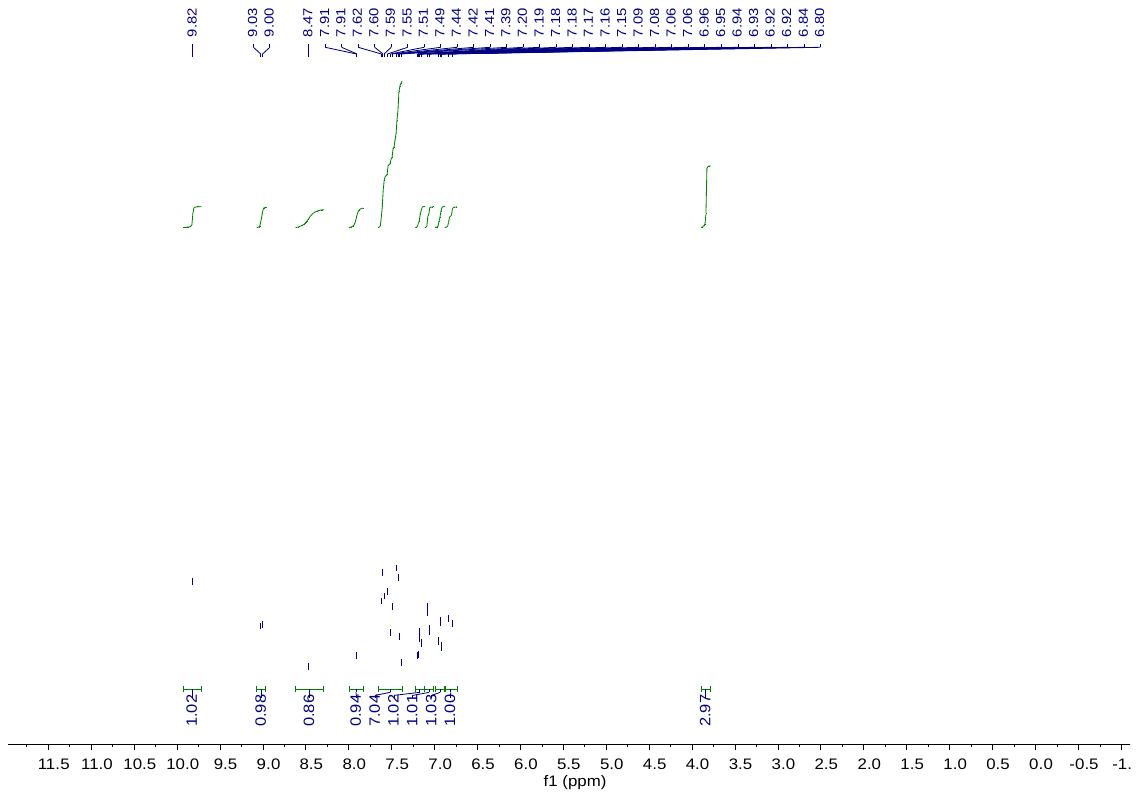
**C2**

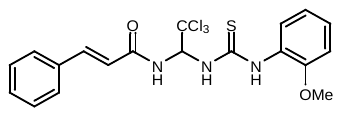

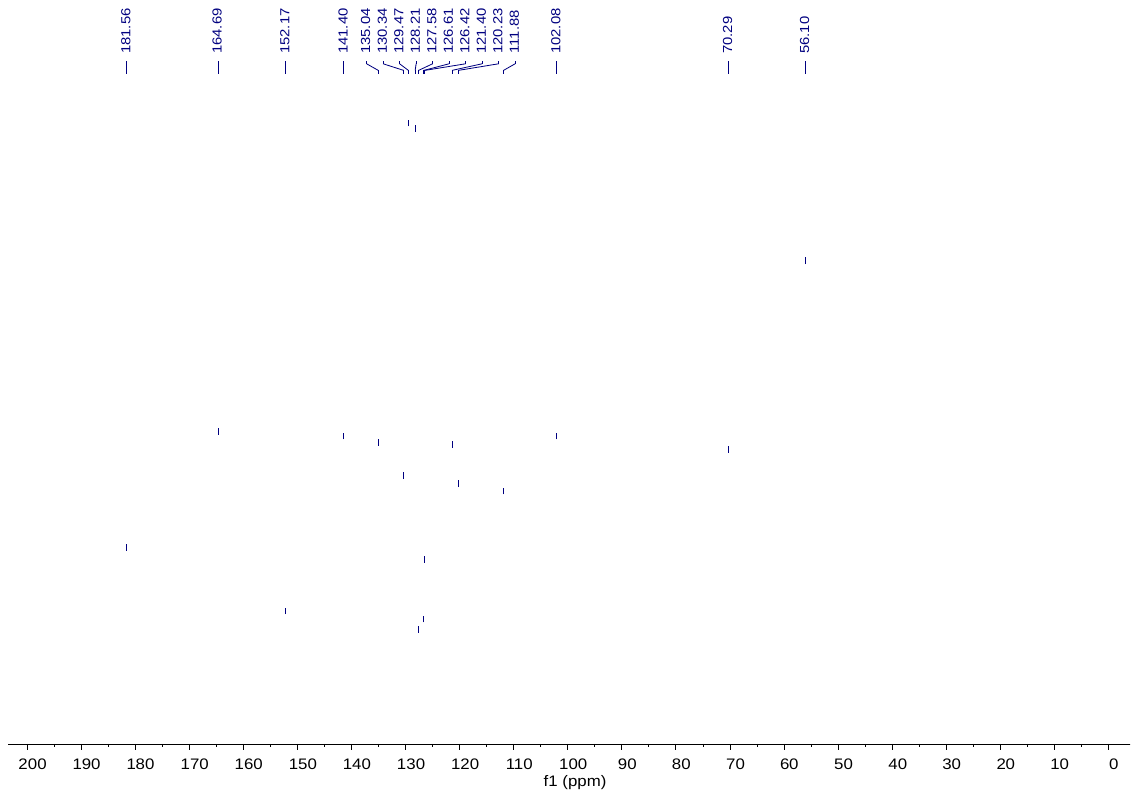
**C2**

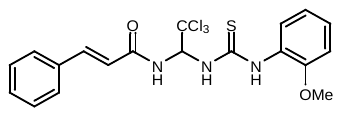

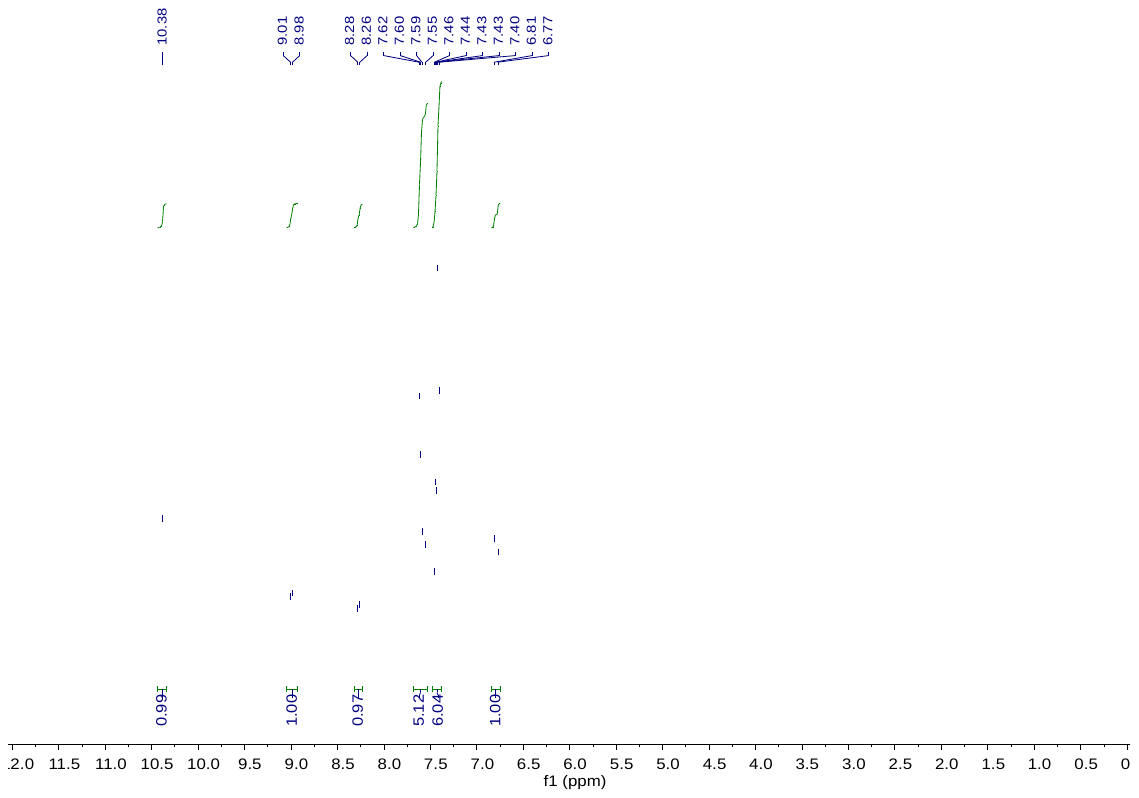

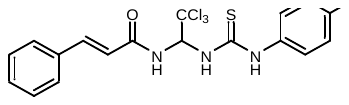
**sal003**

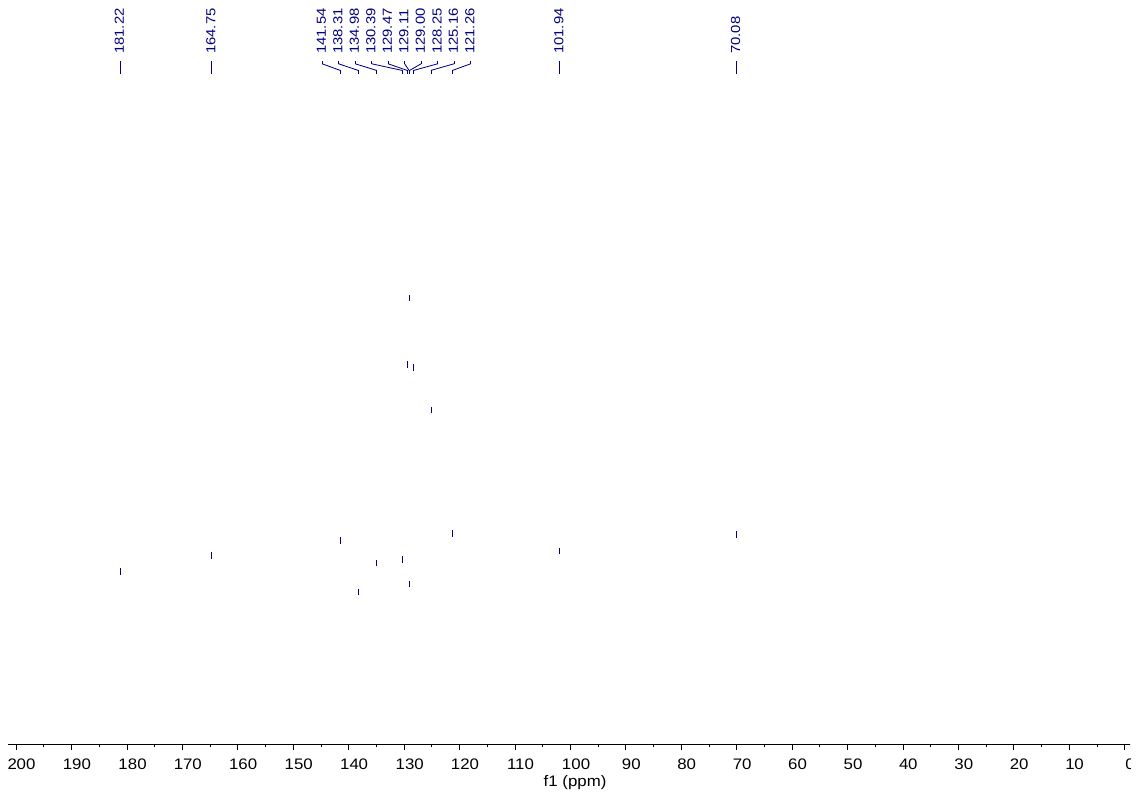

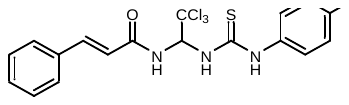
**sal003**

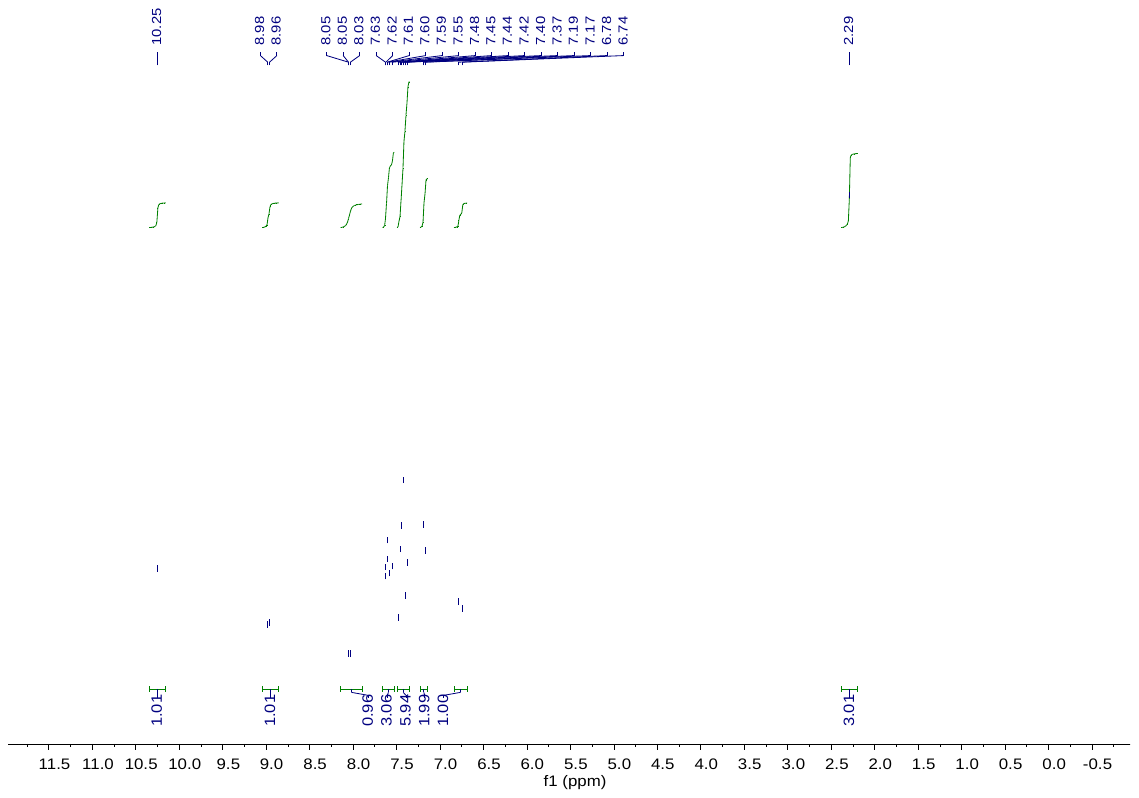

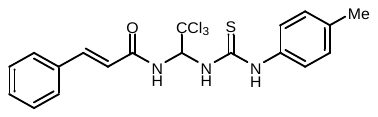
**C4**

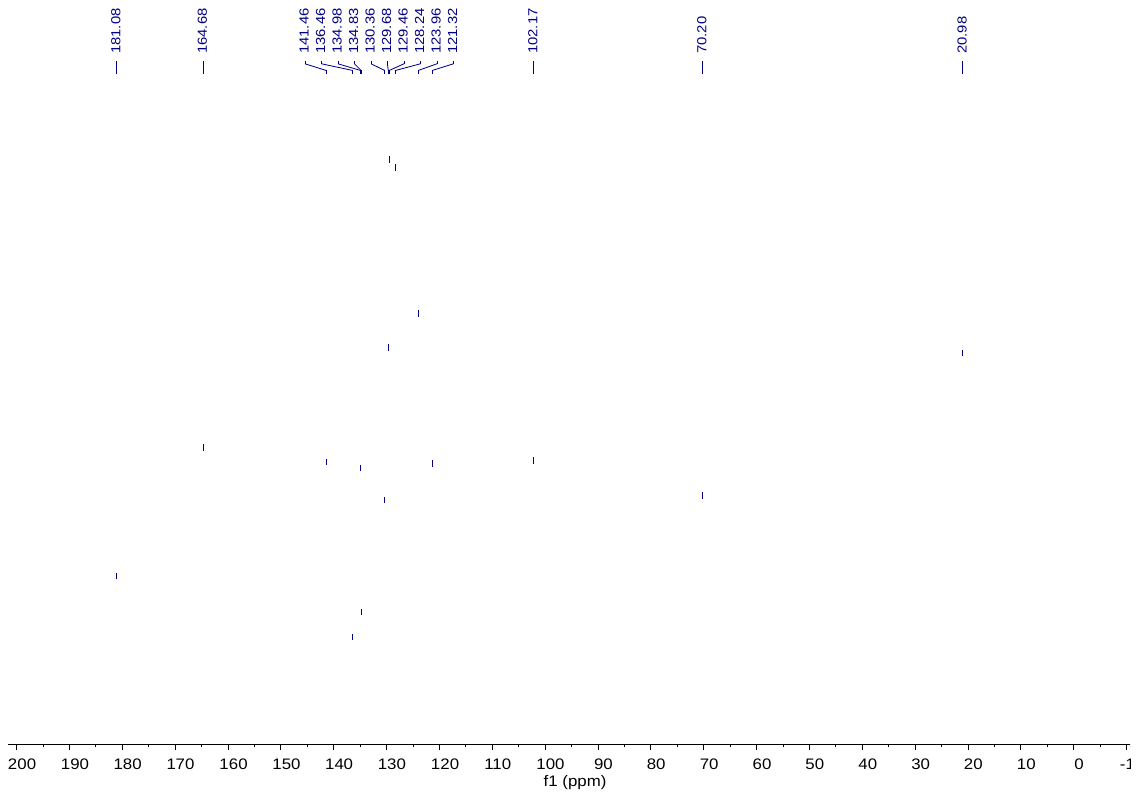
**C4**

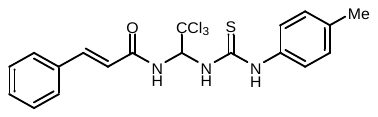

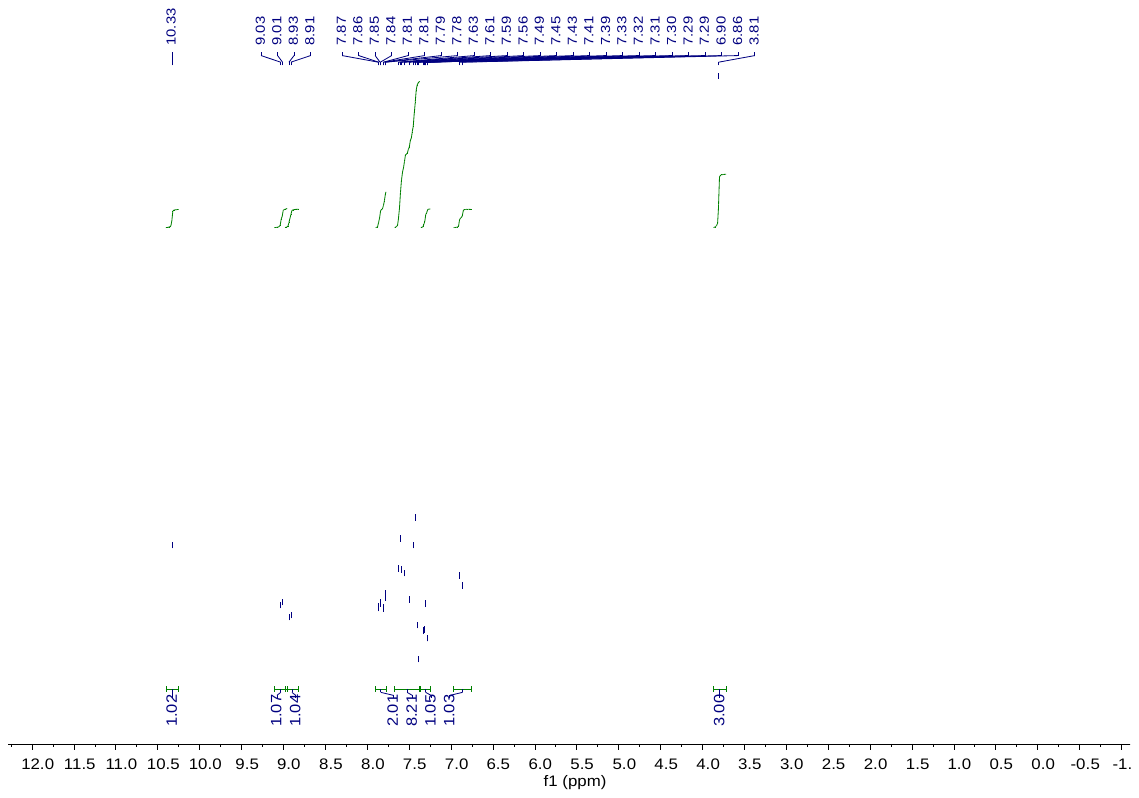
**C5**

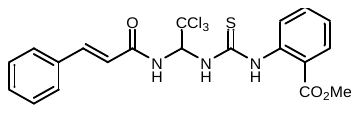

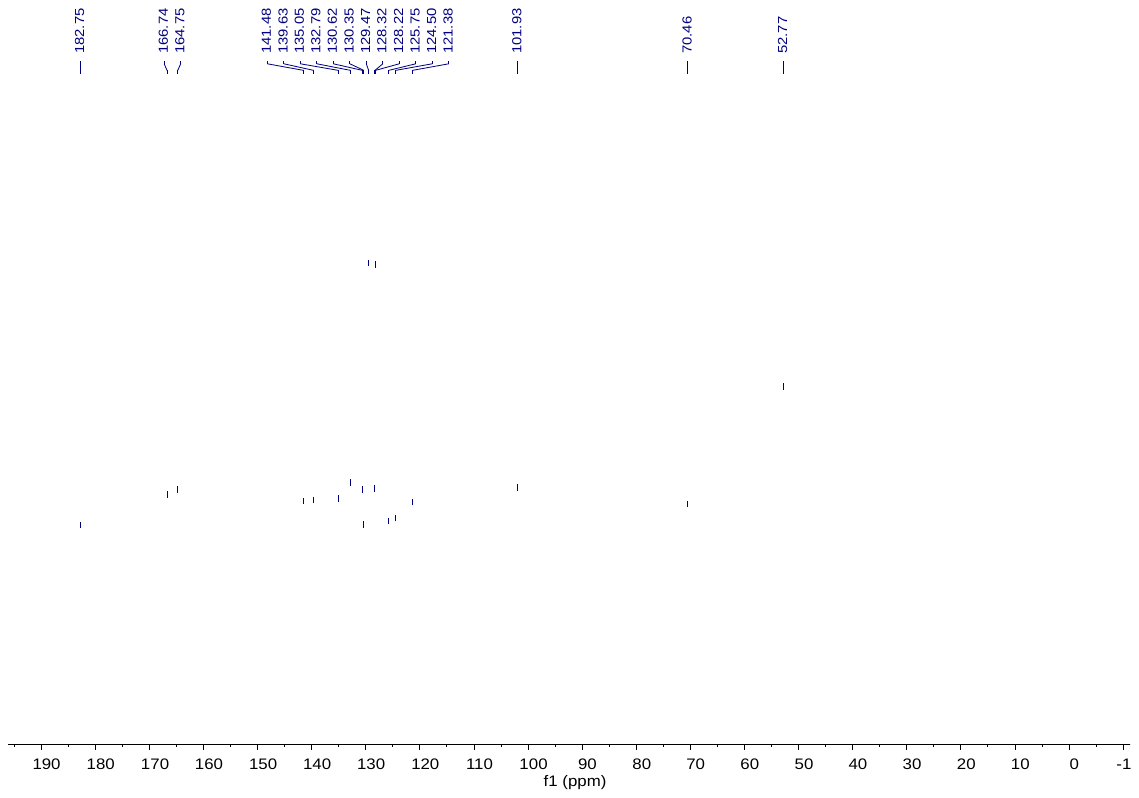
**C5**

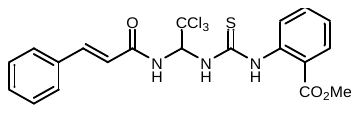

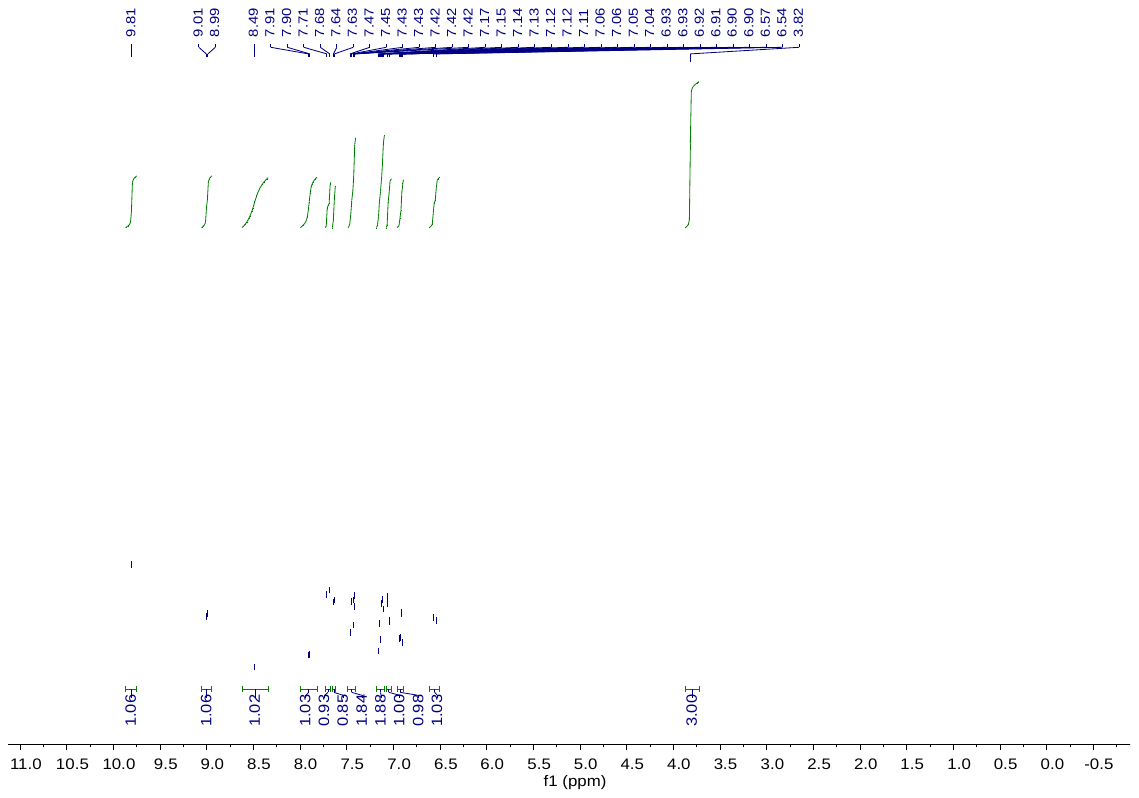

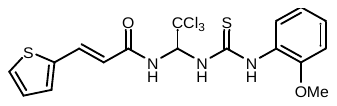
**C6**

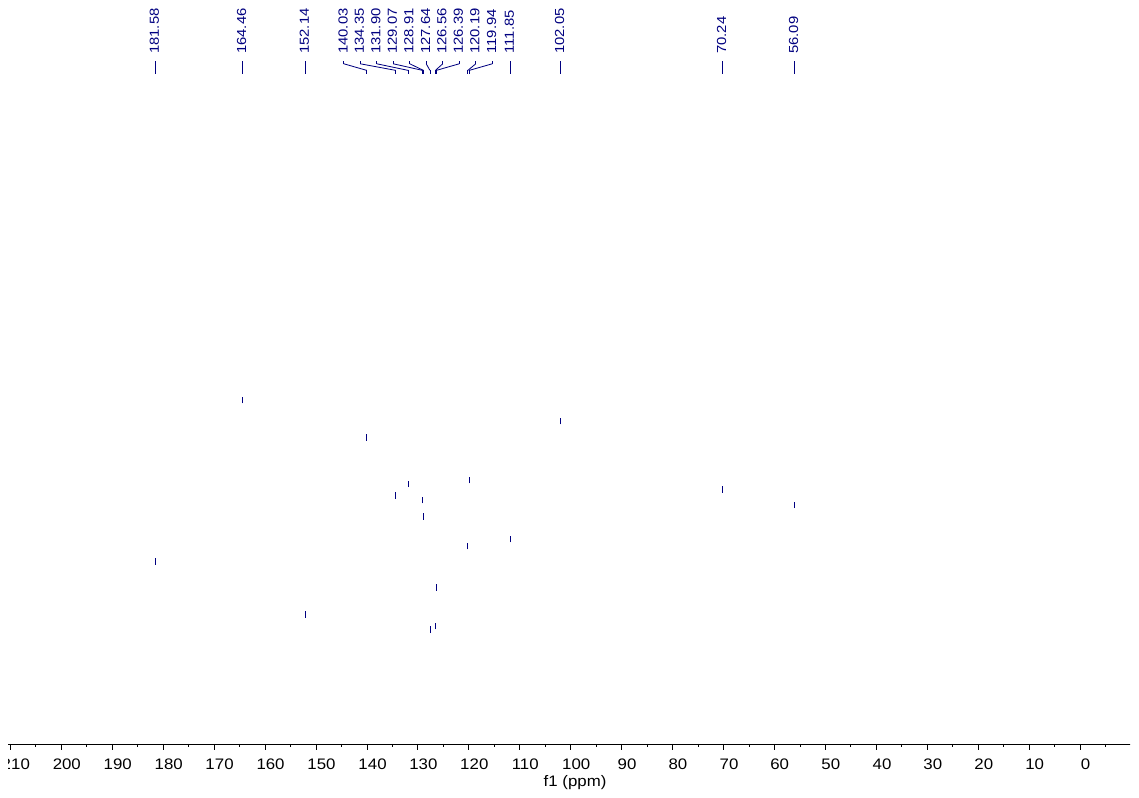

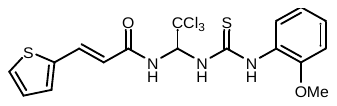
**C6**

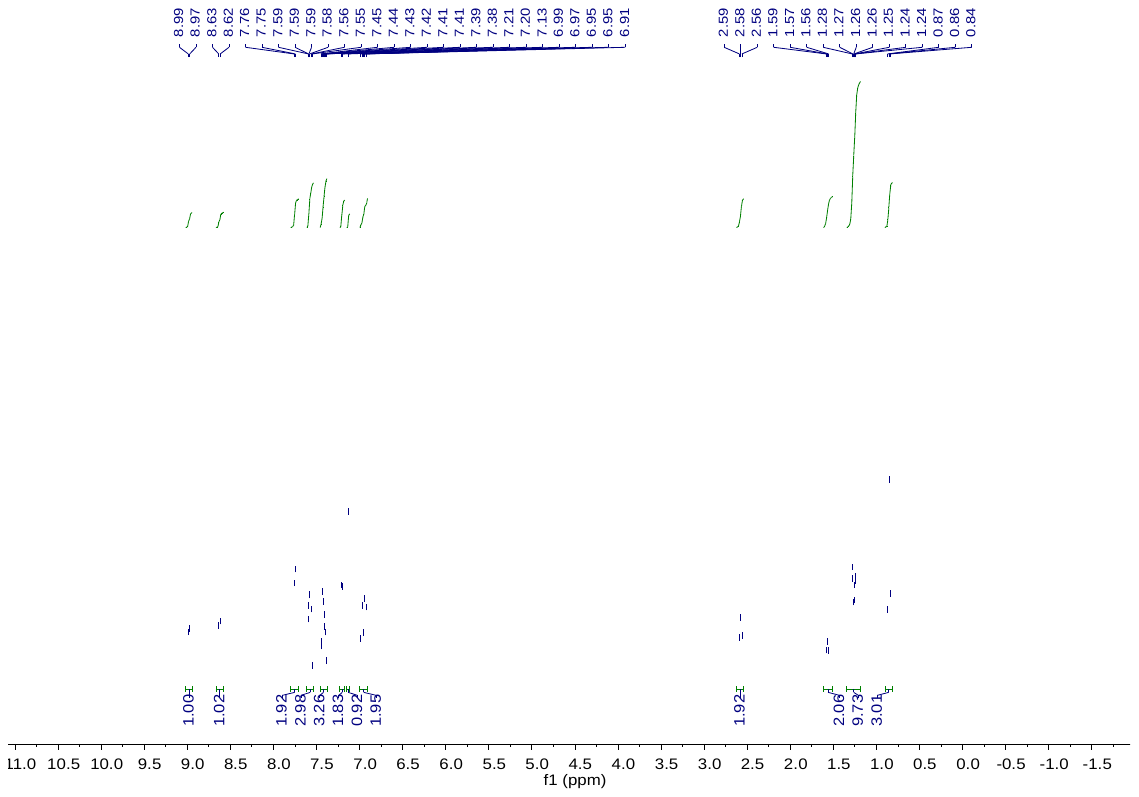
**C7a**

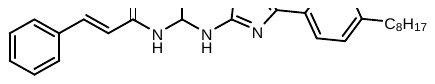

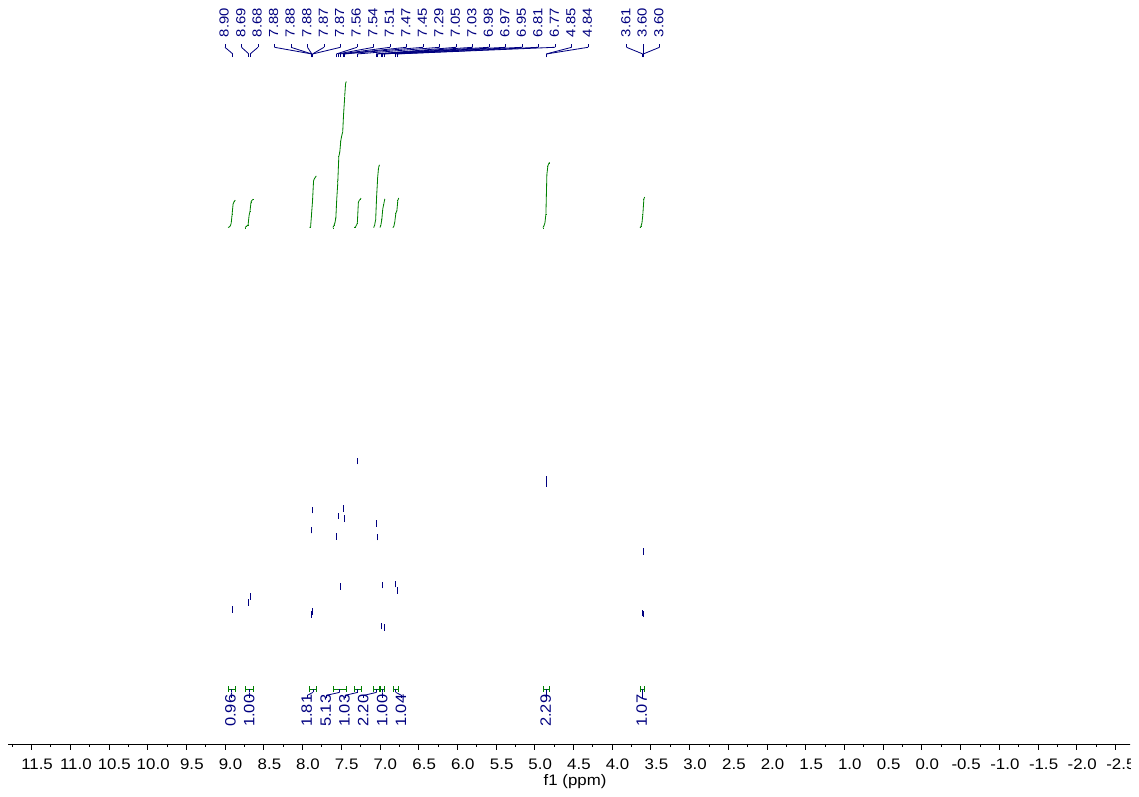
**C7c**

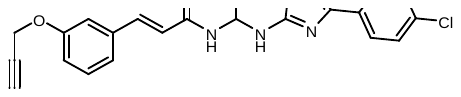

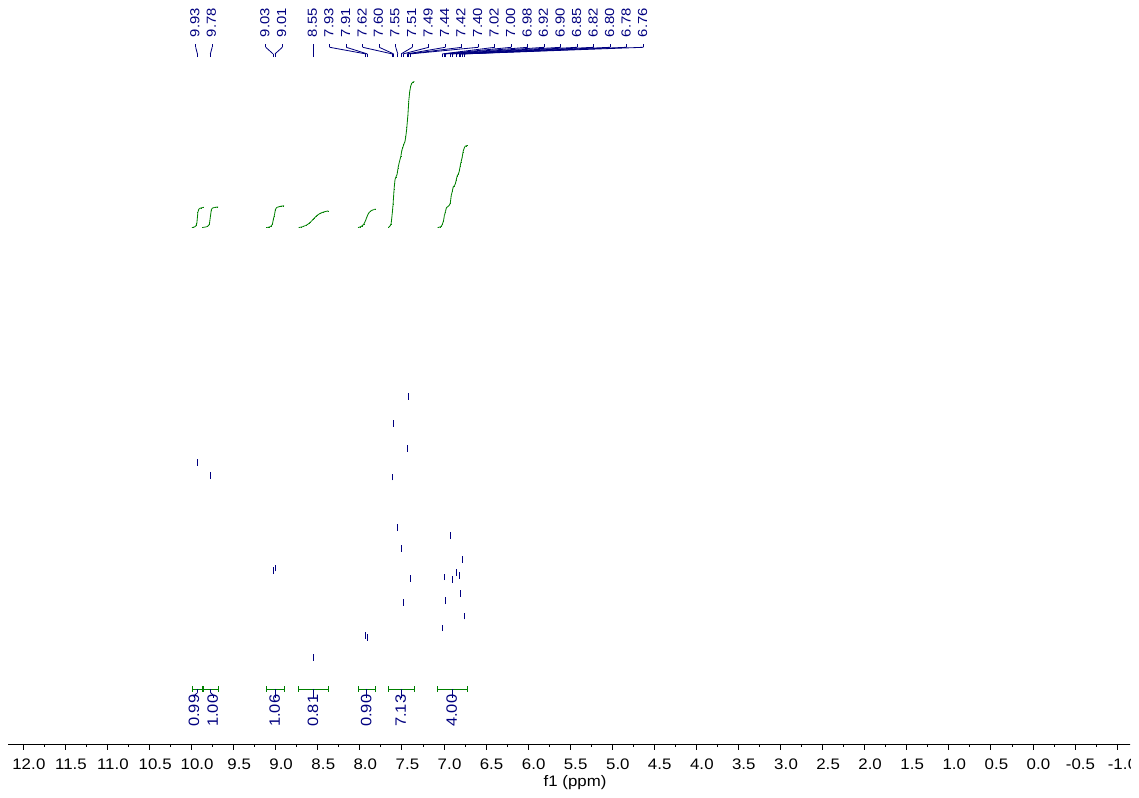

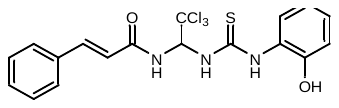
**C9**

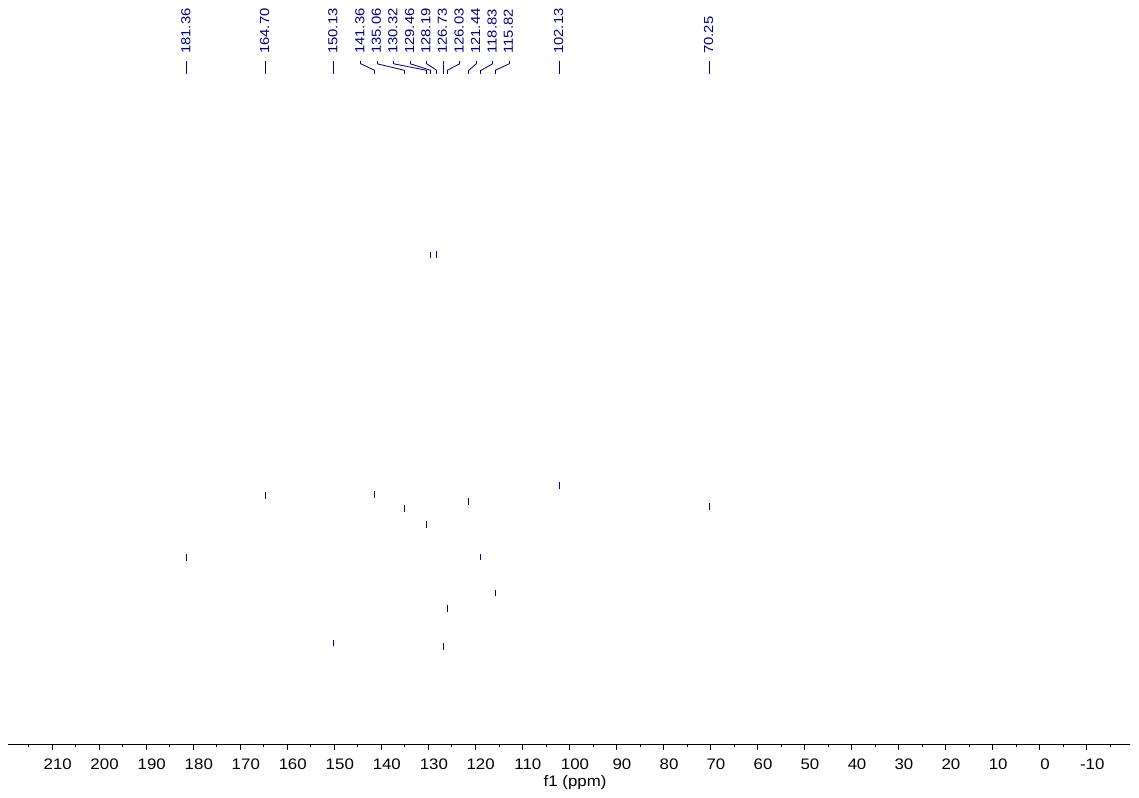
**C9**

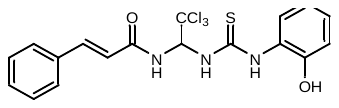

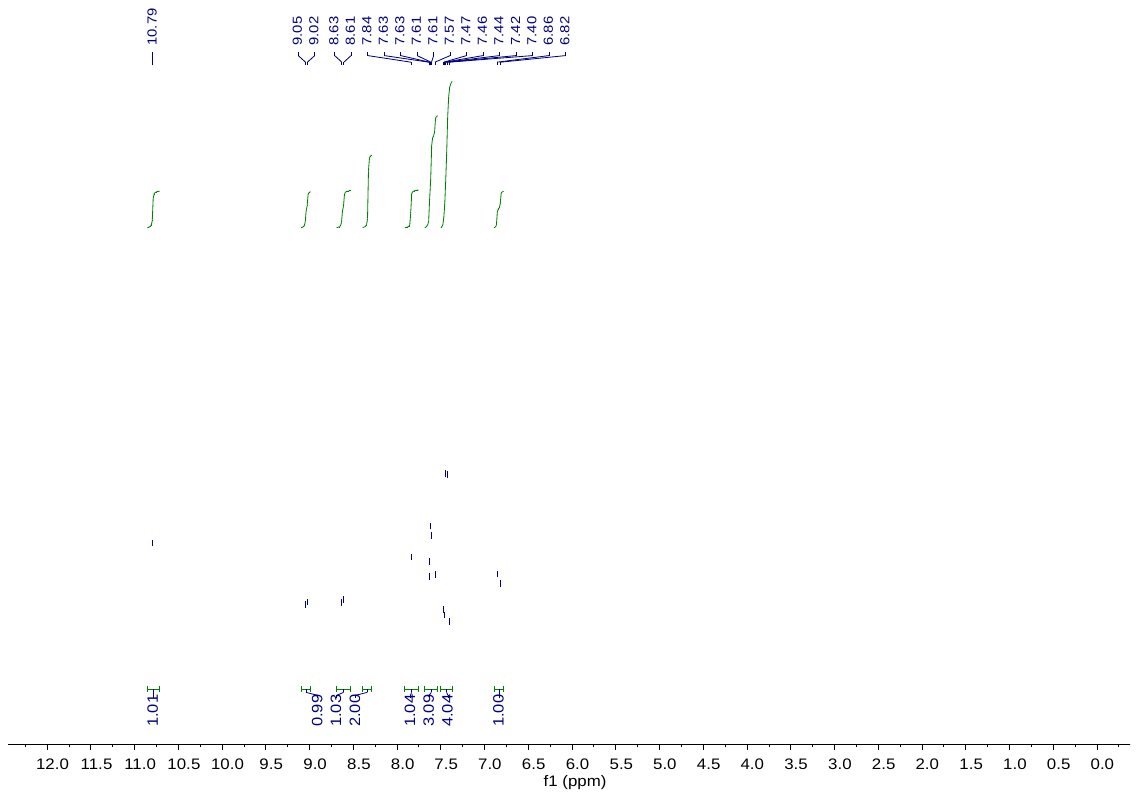

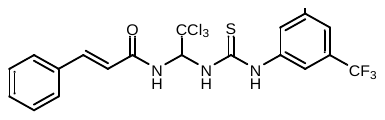
**C10**

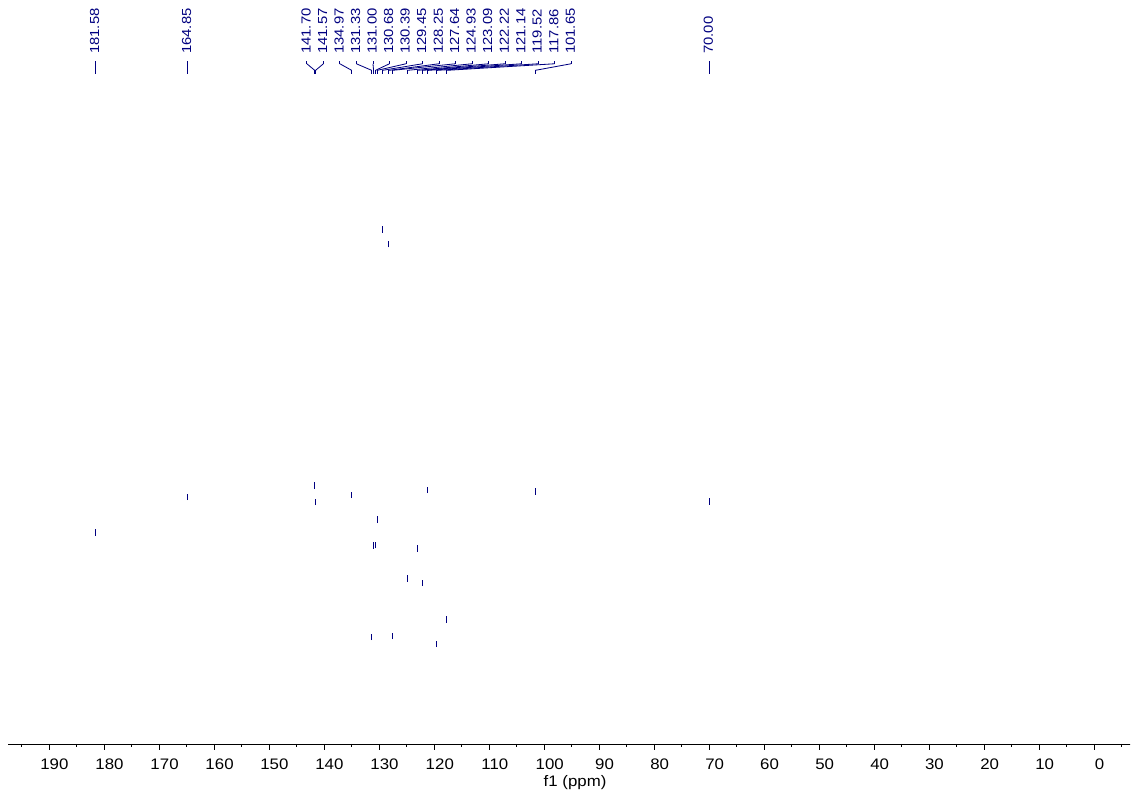
**C10**

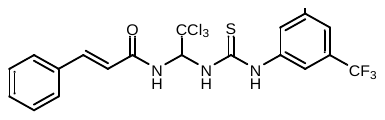

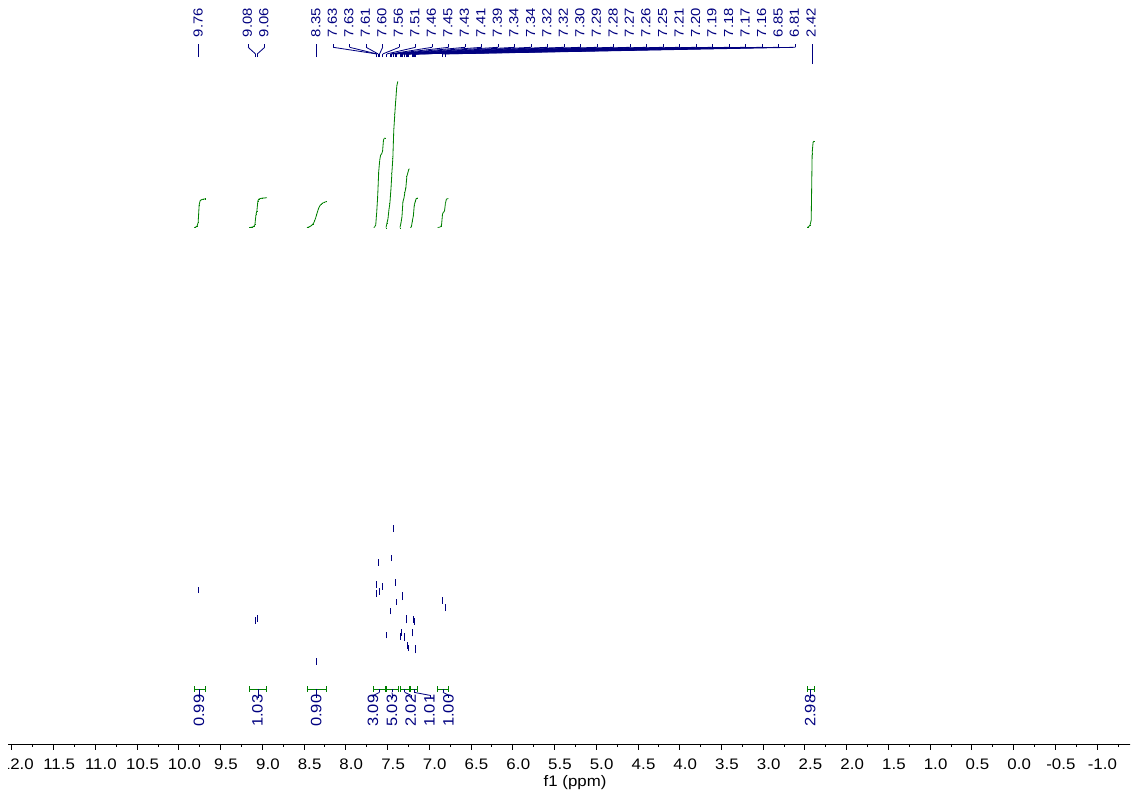
**C11**

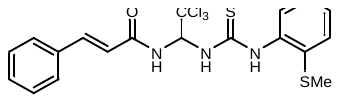

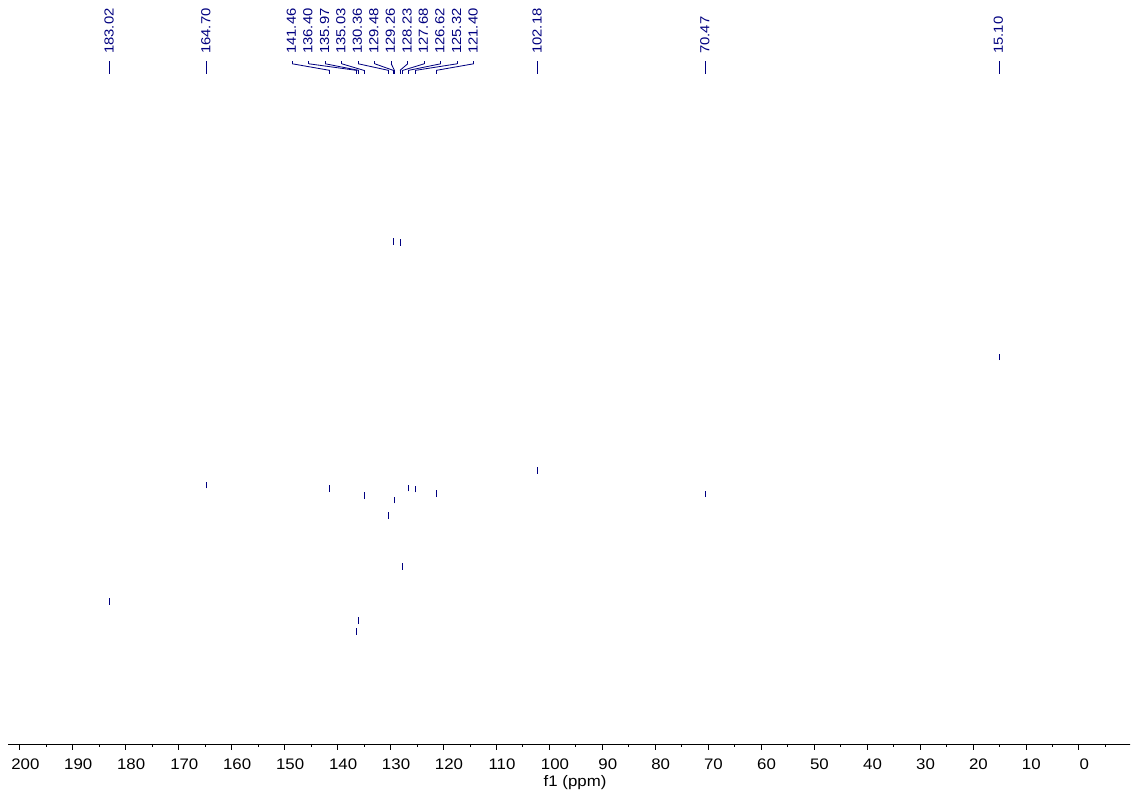
**C11**

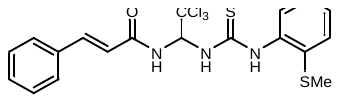

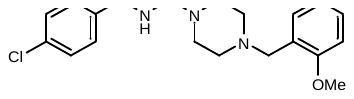

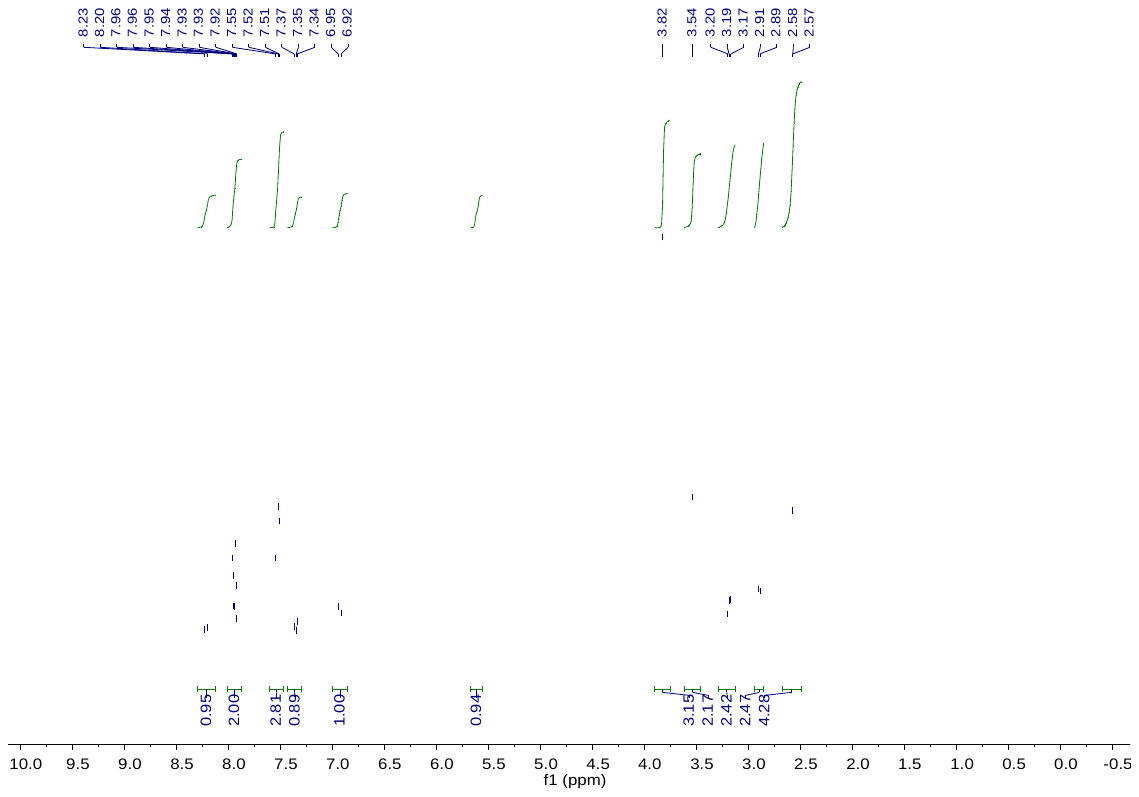
**C13**

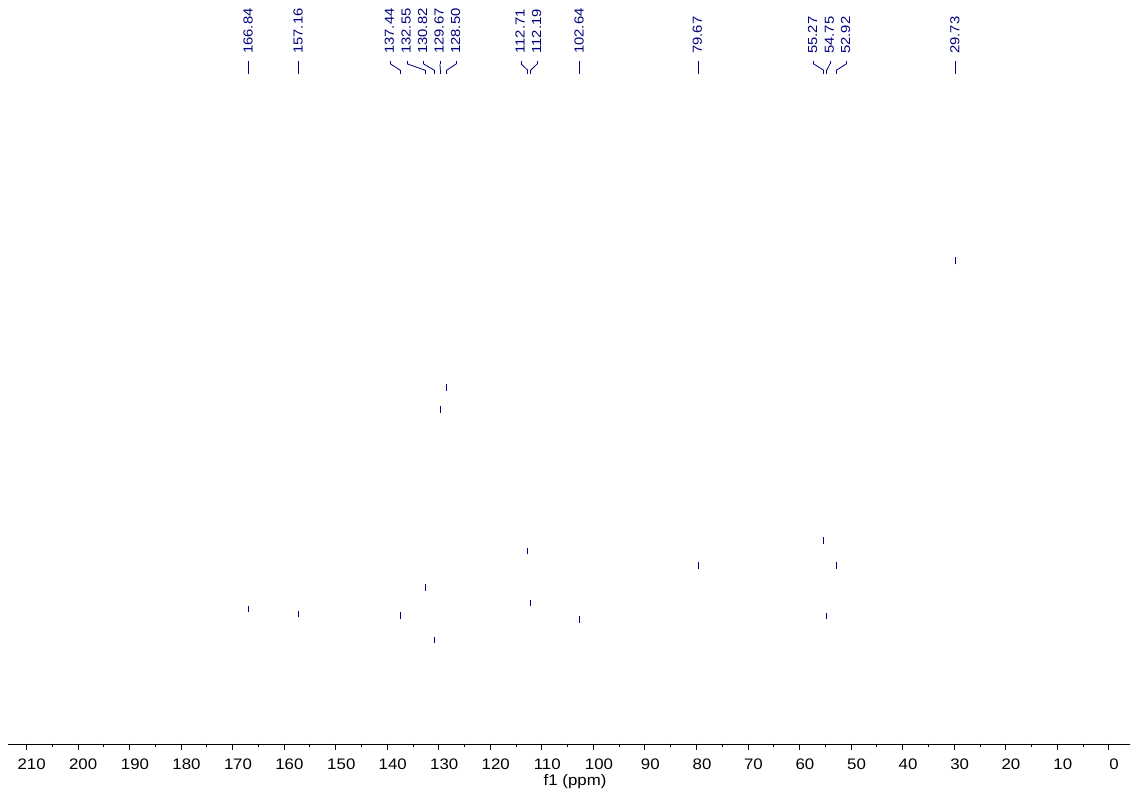

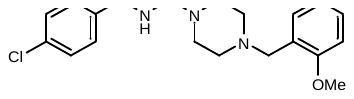
**C13**

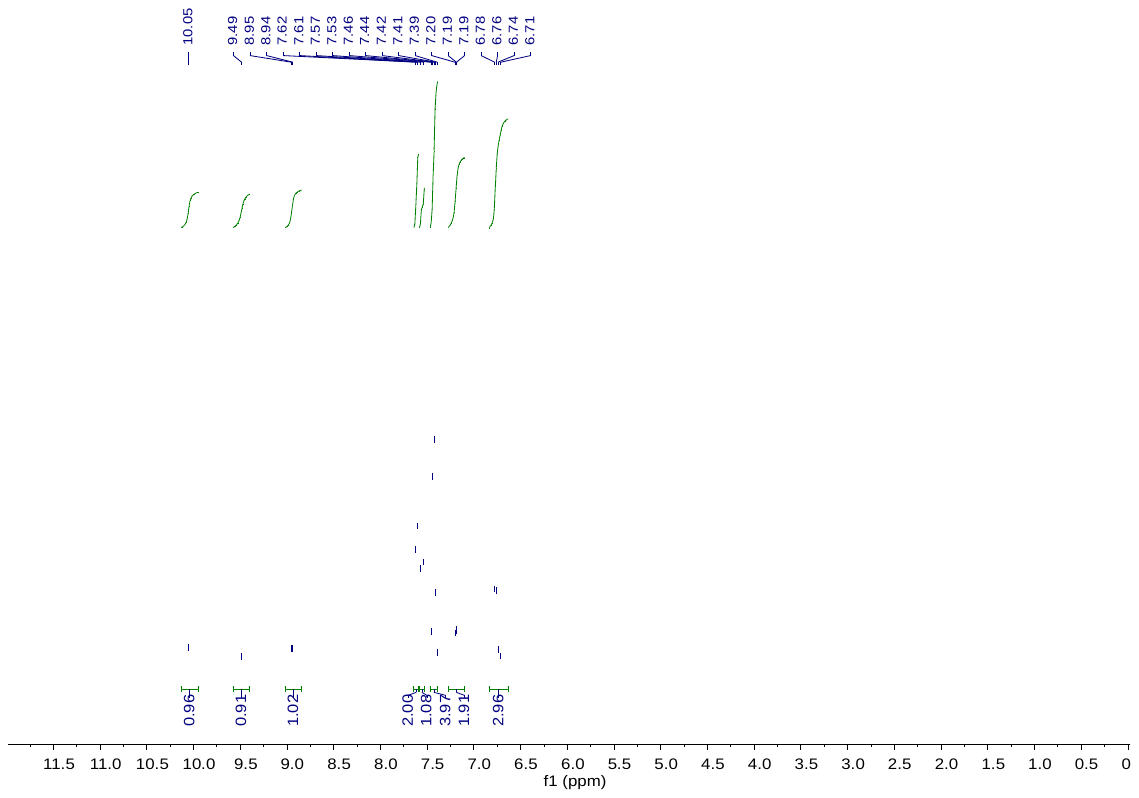
**C14**

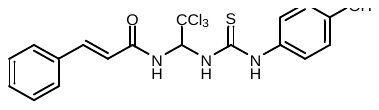

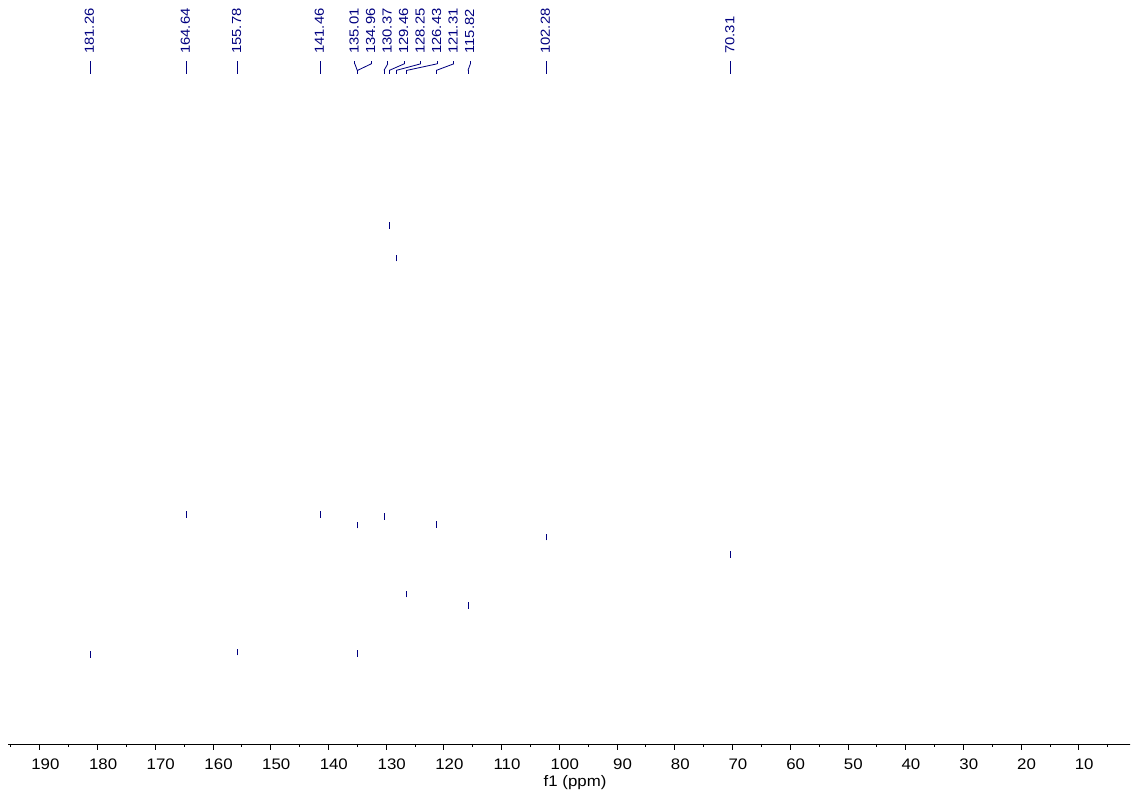
**C14**

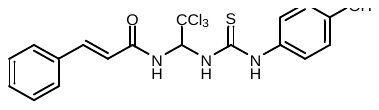

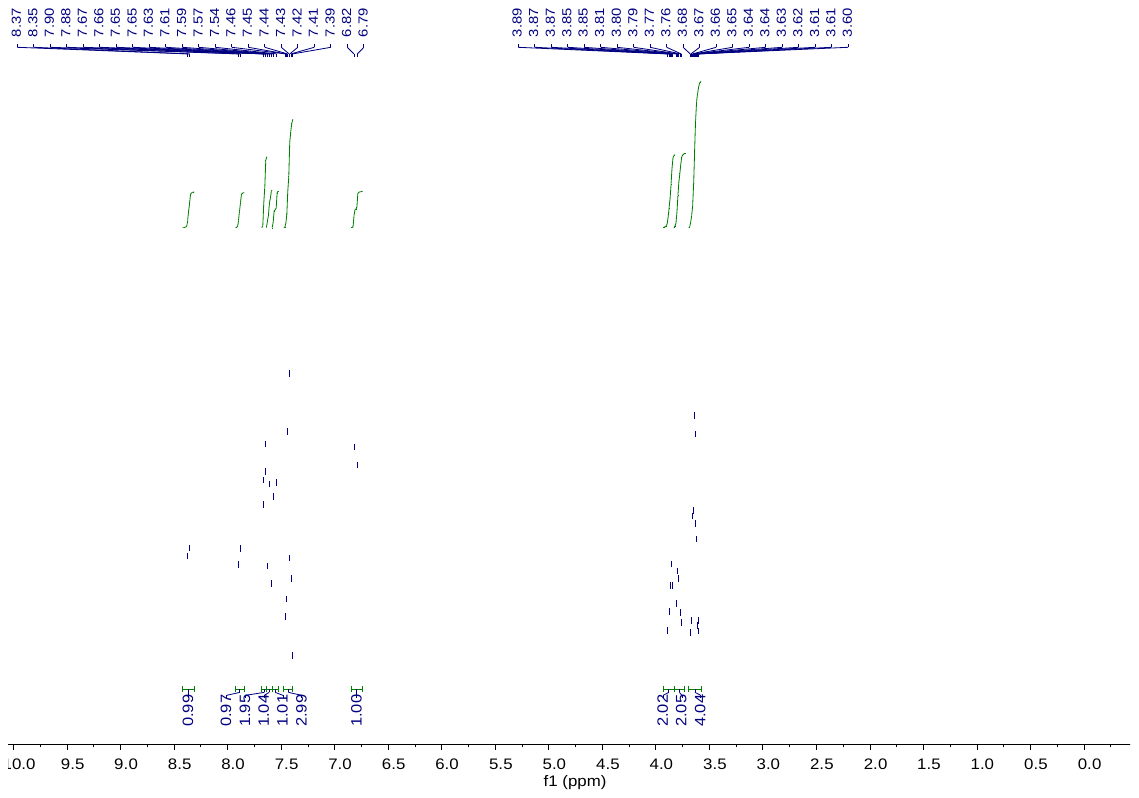
**C16**

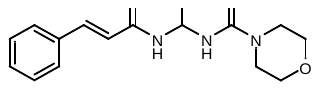

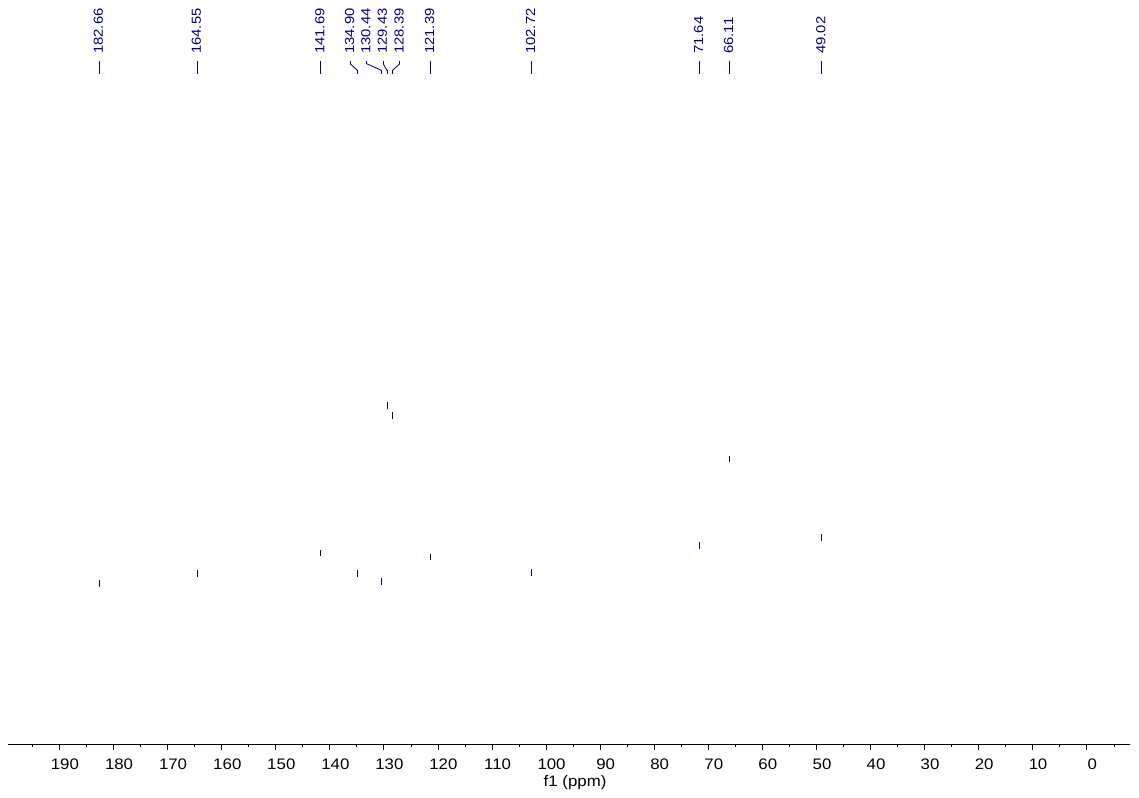
**C16**

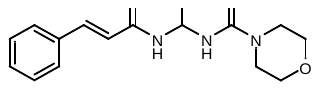

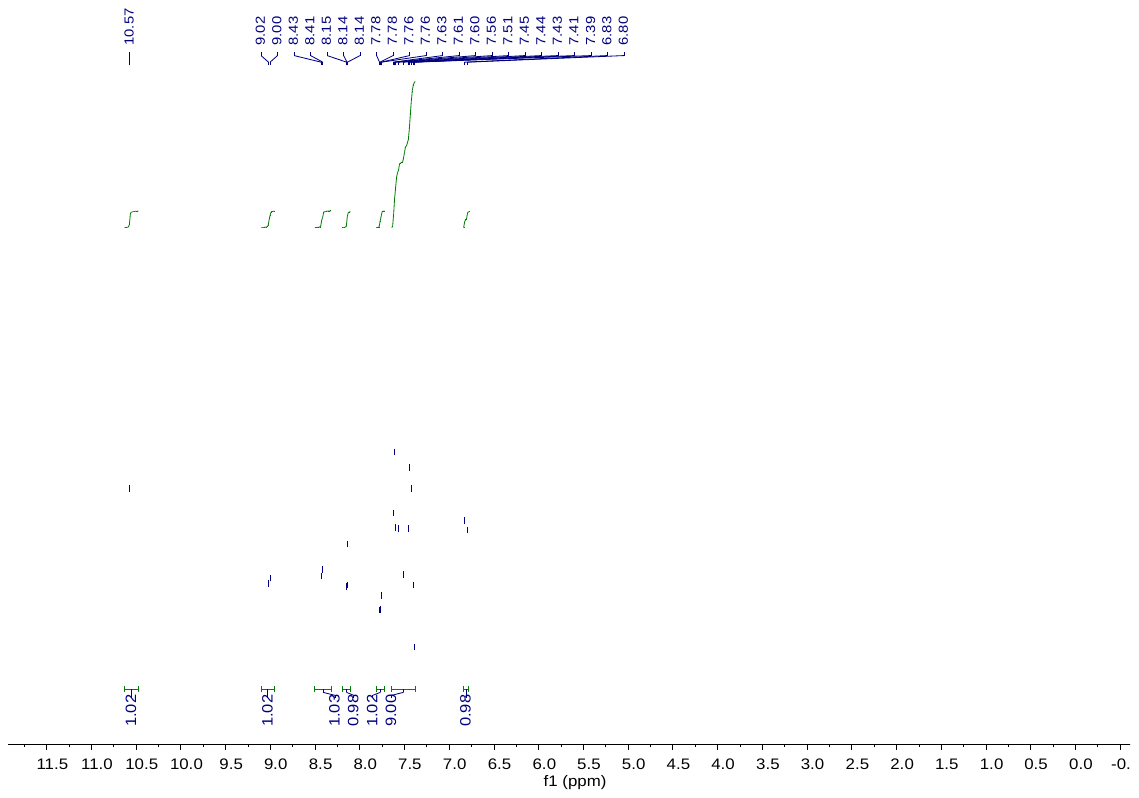

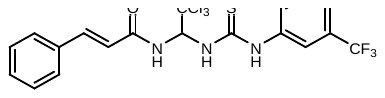
**C17**

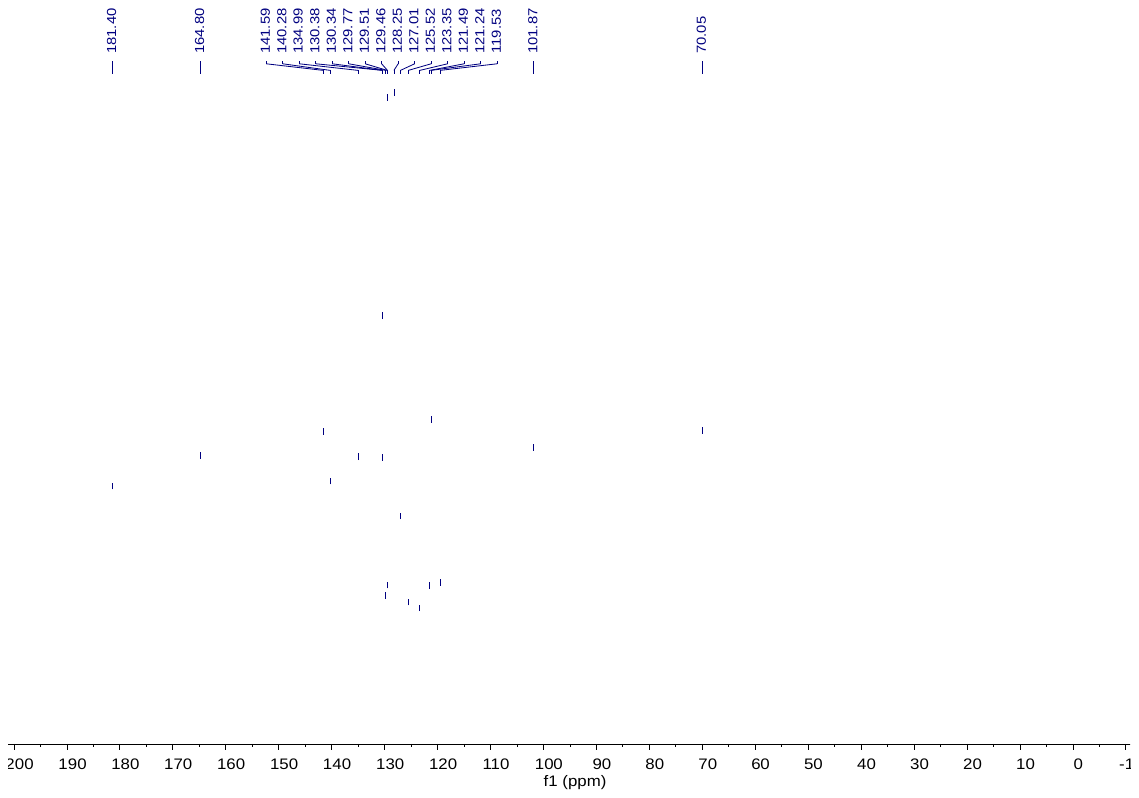
**C17**

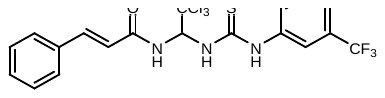

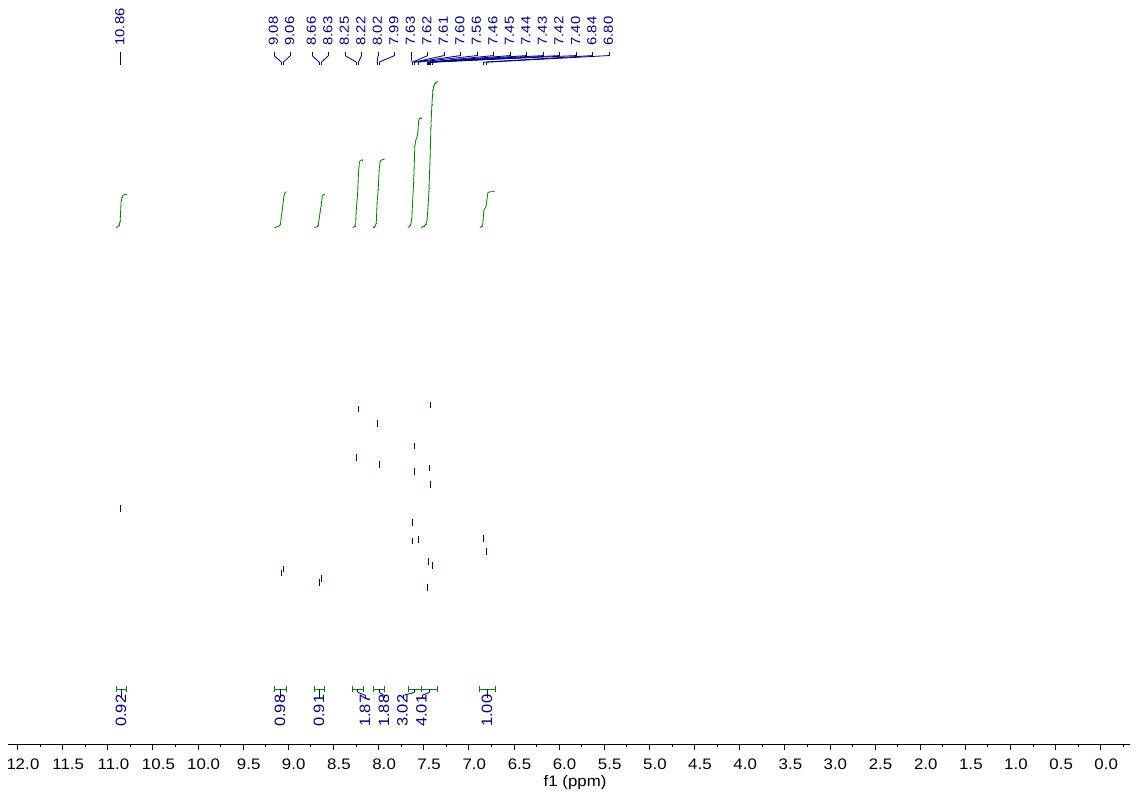

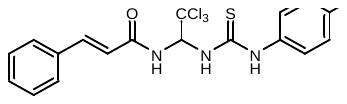
**C18**

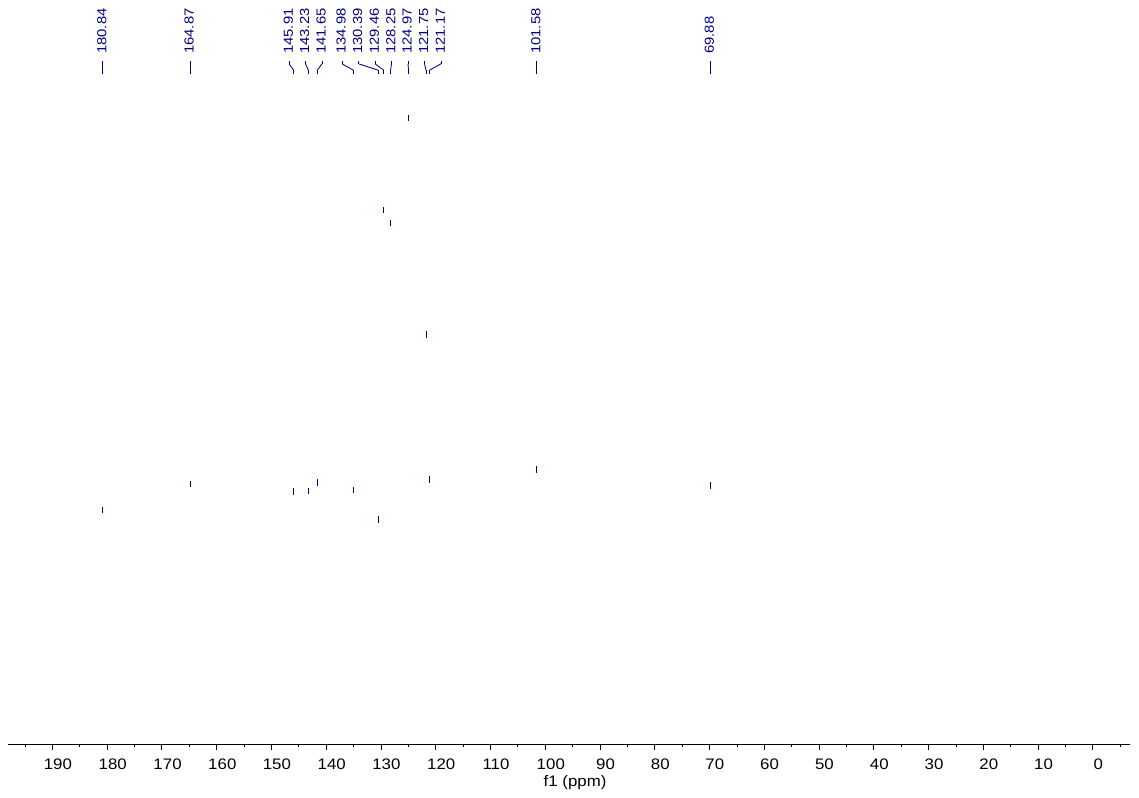
**C18**

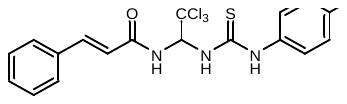

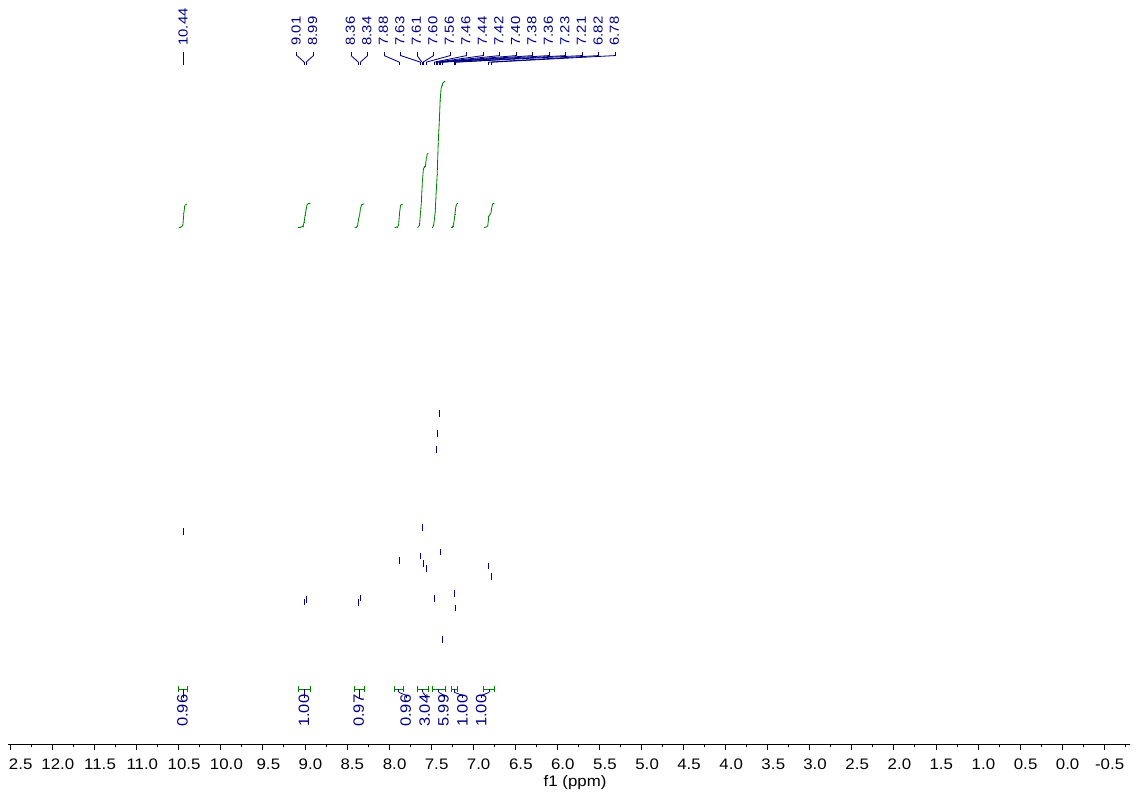

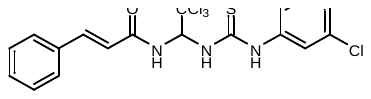
**C19**

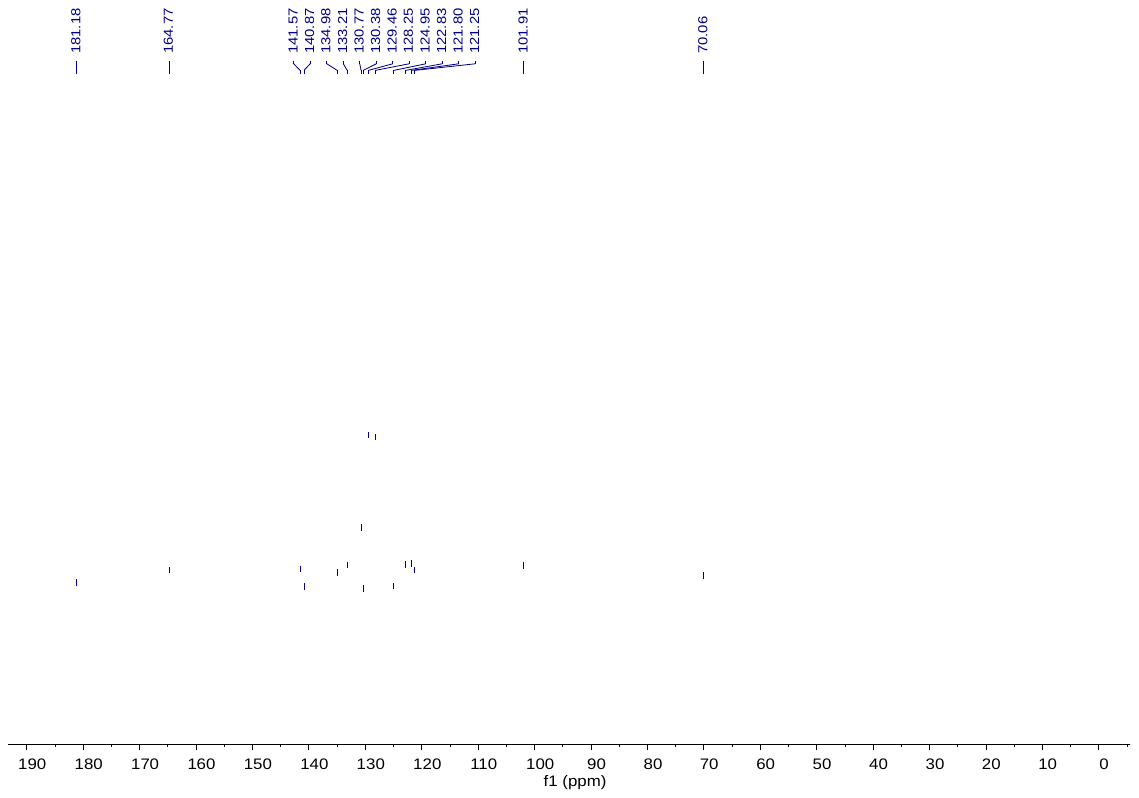
**C19**

**C20**

**C20**
